# Clustering of the structures of protein kinase activation loops: A new nomenclature for active and inactive kinase structures

**DOI:** 10.1101/395723

**Authors:** Vivek Modi, Roland L. Dunbrack

## Abstract

Targeting protein kinases is an important strategy for intervention in cancer. Inhibitors are directed at the active conformation or a variety of inactive conformations. While attempts have been made to classify these conformations, a structurally rigorous catalogue of states has not been achieved. The kinase activation loop is crucial for catalysis and begins with the conserved DFGmotif (Asp-Phe-Gly). This motif is observed in two major classes of conformations, DFGin - a set of active and inactive conformations where the Phe residue is in contact with the C-helix of the N-terminal lobe, and DFGout - an inactive form where Phe occupies the ATP site exposing the C-helix pocket. We have developed a clustering of kinase conformations based on the location of the Phe side chain (DFGin, DFGout, and DFGinter or intermediate) and the backbone dihedral angles of the sequence X-D-F, where X is the residue before the DFGmotif, and the DFG-Phe side-chain rotamer, utilizing a density-based clustering algorithm. We have identified 8 distinct conformations and labeled them based on their Ramachandran regions (A=alpha, B=beta, L=left) and the Phe rotamer (minus, plus, trans). Our clustering divides the DFGin group into six clusters including ‘BLAminus,’ which contains active structures, and two common inactive forms, ‘BLBplus’ and ‘ABAminus.’ DFGout structures are predominantly in the ‘BBAminus’ conformation, which is essentially required for binding Type II inhibitors. The inactive conformations have specific features that make them unable to bind ATP, magnesium ion, and/or substrates. Our structurally intuitive nomenclature will aid in understanding the conformational dynamics of these proteins and structure-based development of kinase drugs.

**Significance statement:** Protein kinases play important roles in signaling pathways and are widely studied as drug targets. Their active site exhibits remarkable structural variation as observed in the large number of available crystal structures. We have developed a clustering scheme and nomenclature to categorize and label all the observed conformations in human protein kinases. This has enabled us to clearly define the geometry of the active state and to distinguish closely related inactive states which were previously not characterized. Our classification of kinase conformations will help in better understanding the conformational dynamics of these proteins and the development of inhibitors against them.

## INTRODUCTION

Phosphorylation is a fundamental mechanism by which signaling pathways are regulated in cells. Protein kinases are cellular sentinels which catalyze the phosphorylation reaction by transferring the γ-phosphate of an ATP molecule to Ser, Thr, or Tyr residues of the substrate. Due to their crucial role in the functioning of the cell, protein kinases are tightly regulated. Dysregulation of kinases may result in variety of disorders including cancer, making development of compounds for modulating kinase activity an important therapeutic strategy.

The human genome contains ~500 protein kinases that share a common fold consisting of two lobes: an N-terminal lobe, consisting of a five stranded β-sheet with an α-helix called the C-helix, and a C-terminal lobe comprising six α-helices (Fig. 1). They are divided broadly into nine families based on their sequences (1). The two lobes are connected by a flexible hinge region forming the ATP binding site in the middle of the protein. The active site comprises several structural elements that are crucial for enzymatic activity. The activation loop is typically 20-30 residues in length beginning with a conserved DFG motif (usually Asp-Phe-Gly)and extending up to an APE motif (usually Ala-Pro-Glu). In active kinase structures, this loop forms a cleft that binds substrate. Bound substrate peptide forms specific interactions with the conserved HRD motif (usually His-Arg-Asp) which occurs in the catalytic loop of the protein. In the active conformation, Asp is in a position and orientation to bind a magnesium ion that interacts directly with an oxygen atom of the β phosphate of ATP. The active state exhibits an inward disposition of the C-helix and a salt bridge interaction between a conserved Lys residue in the β3 strand and a Glu residue in the C-helix. The lysine side chain forms hydrogen bonds with oxygen atoms of the α and β phosphates of ATP. The N-lobe has a GxGxxG motif in a loop that stabilizes the phosphates of the bound ATP molecule during catalysis. The regulation of the activity of a kinase is achieved in part by the plasticity of these elements of the structure (2). The catalytically active state of a kinase requires a unique assembly of these elements that create an environment conducive to the phosphotransfer reaction.

**Figure 1.**
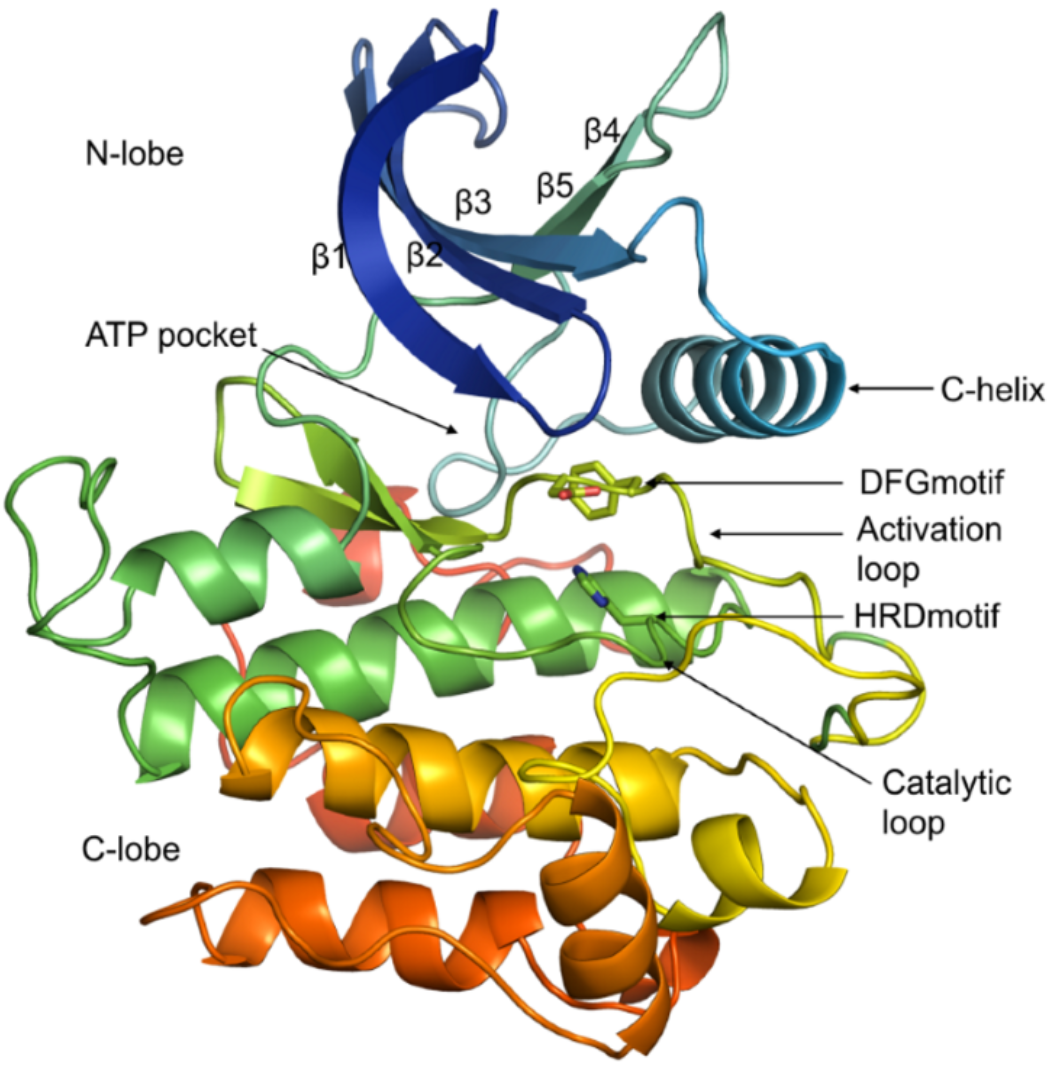
Structure of a typical protein kinase domain displaying ATP binding site and conserved elements around it (INSR kinase, PDB id 1GAG).

Inactive states of a kinase do not have the chemical constraints required for catalytic activity and therefore kinases exhibit multiple inactive conformations (3). Typically, in an inactive conformation the activation loop is collapsed onto the surface of the protein, blocking substrate binding and rendering the kinase catalytically inactive. In addition, many inactive conformations have positions of the DFGmotif incompatible with binding ATP and magnesium ion required for catalysis. In the DFGout conformation, DFG-Phe and DFG-Asp swap positions so that DFG-Phe occupies the ATP binding pocket and DFG-Asp is *out* of the active site. There are diverse DFGin structures from multiple kinases where DFG-Phe remains adjacent to the C-helix but in a different orientation (and sometimes position) from that of active DFGin structures. There are also structures where the Phe is in positions intermediate between the typical DFGin and DFGout states. The many inactive, non-DFGout conformations have been variously referred to as pseudo DFGout, DFGup, SRC-like inactive, and atypical DFGout (4, 5). Although, DFGin and DFGout are broadly recognized groups of conformations, a consensus nomenclature for the inactive states is lacking.

The DFGin and DFGout conformations have been used as the basis of grouping the inhibitors developed against the active site of these proteins into two main categories (6, 7). Molecules such dasatinib which occupy the ATP pocket only are called Type I inhibitors, and typically bind DFGin conformations but not exclusively. Type II Inhibitors like imatinib bind to the DFGout state and extend into the hydrophobic allosteric pocket underneath the C-helix (8). Design of better inhibitors could be guided by a better understanding and classification of the conformational variation observed in kinases.

The number of mammalian kinase structures in the Protein Data Bank (PDB) has risen to ~3300 entries from ~250 kinases capturing the remarkable conformational diversity of this protein family. There have been some attempts to classify kinase conformations and to study inhibitor interactions (9–12). Möbitz has performed a quantitative classification of all the mammalian kinases using pseudo dihedral angles of four consecutive Cα atoms of the residues of the DFGmotif and its neighbors and its distance from the C-helix (11). This resulted in a scheme dividing kinase conformations into twelve categories with labels such as ‘FG-down,’ ‘FG-down αC-out,’ ‘G-down αC-out,’ ‘A-under P BRAF,’ ‘A-under P-IGF1R,’ etc. Recently, Ung and coworkers used a similar idea of using two directional vectors for the DFGmotif residues and the distance from the C-helix to classify kinases into five groups, C-helix-in-DFGin (CIDI), C-helix-in-DFGout (CIDO), C-helix-out-DFGin (CODI), C-helix-out-DFGout (CODO), and ωCD (12). Some other classification schemes have emphasized the binding modes of inhibitors (4, 13).

In this paper, we present a new clustering and classification of the conformational states of protein kinases that addresses some of the deficiencies of previous such efforts. These deficiencies include either too few or too many structural categories, failing to distinguish DFGin inactive conformations from active structures, and an inability to automatically classify new structures added to the PDB. In the current work, we have clustered all the human kinase structures at two levels of structural detail. First, at a broader level we grouped kinase structures into three categories depending on the spatial position of the DFG-Phe side chain. These three groups are labeled the DFGin, DFGout, and DFGinter (intermediate) conformations. Second, we clustered each of the three spatial groups at a finer level based on the dihedral angles required to place the Phe side chain: the backbone dihedral angles ϕ and ψ of the residue preceding the DFGmotif (X-DFG), the DFG-Asp residue, and the DFG-Phe residue, as well as the χ_1_side-chain dihedral angle of the DFG-Phe residue. This produced a total of eight clusters - six for DFGin, and one cluster each for the DFGout and DFGinter groups.

We have developed a nomenclature that is intuitive to structural biologists based on the regions of the Ramachandran map occupied by the X,D, and F residues of the X-DFG motif (‘A’ for alpha helical region, ‘B’ for beta sheet region, ‘Ľ for left-handed helical region) and the χ_1_ rotamer of the Phe side chain (‘minus’ for the −60° rotamer; ‘plus’ for the +60° rotamer; and ‘trans’ for the 180° rotamer). We have clearly defined the active state of kinases, designated ‘BLAminus,’ which is the most common kinase conformation in the PDB. Further, we also clearly define different inactive DFGin conformations which were previously grouped together. The most common inactive DFGin conformations are BLBplus and ABAminus. The Type II-binding DFGout state is labeled BBAminus. Overall, our clustering and nomenclature scheme provides a structural catalogue of human kinase conformations which will provide deeper insight into the structural variation of these proteins, benefitting structure-guided drug design.

## RESULTS

### Clustering kinase conformations based on spatial location of the DFG-Phe residue

Human protein kinases in the PDB were identified by sequence, excluding proteins such as PI3-PI4 kinases that are distantly related to canonical protein kinases but possess highly divergent folds. This led to a dataset with 244 human kinase domain sequences with known structures from 3,343 PDB entries, having 4,834 polypeptide chains containing kinase domains (some asymmetric units have multiple copies of the kinase). To identify a set of reliable structures where the catalytic machinery is primed to catalyze the phosphorylation reaction and the activation loop orientation is conducive to substrate binding we selected structures that satisfy the following criteria: 1) resolution ≤ 2.25 Å and R-factor ≤ 0.25; 2) ATP or a triphosphate analogue of ATP bound to the active site (PDB ligand codes ATP, ANP, ACP or AGS); 3) Mg^2+^ or Mn^2+^ ion bound in the active site; and 4) a phosphorylated Ser, Thr or Tyr residue in the activation loop. Not all kinases require a phosphorylated residue in the activation loop for activity, but its presence is usually indicative of an active kinase structure. This led to the identification of a set of 28 chains from 24 PDB entries and 12 different kinases (listed in Table S1). We refer to these structures as ‘catalytically primed.’ An example is shown in Fig. 2A. Dihedral angle features of these structures are plotted in Fig. 2B and Fig. 2C and discussed further below.

**Figure 2.**
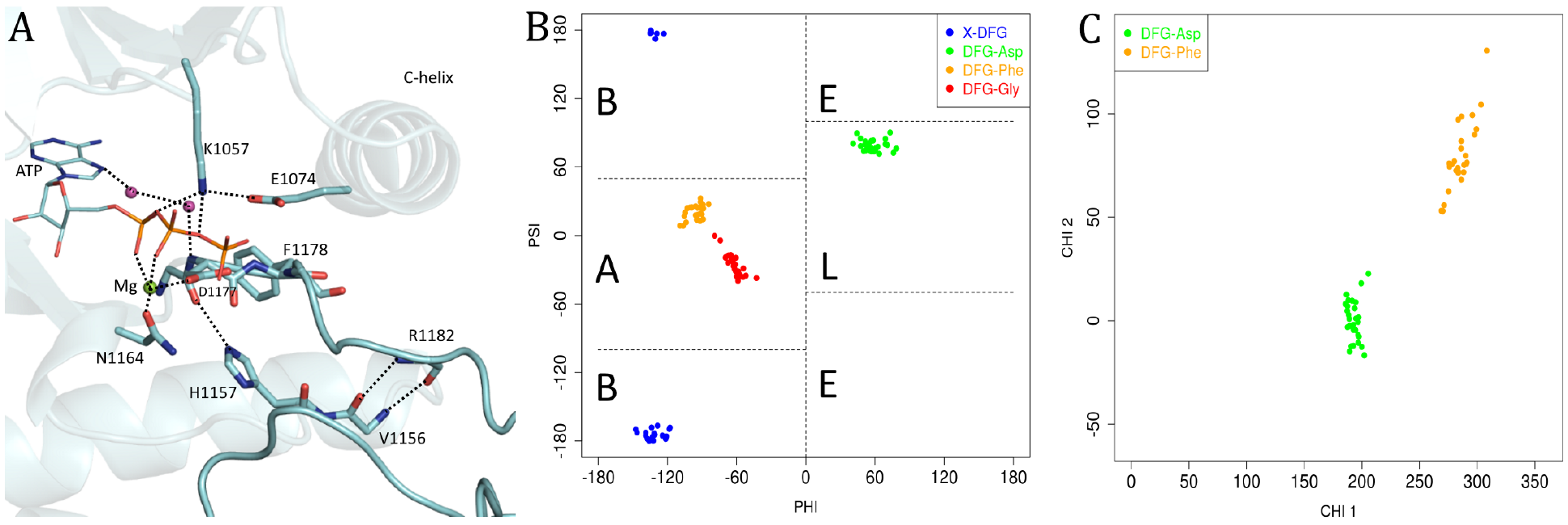
Analysis of catalytically primed structures. (*A*) Catalytically primed structure of insulin receptor kinase (PDB: 3BU5 chain A) with bound ATP and Mg ion. Important interactions are marked with dashed lines. (*B*) Ramachandran plot of catalytically primed structures (those with bound ATP and Mg, a phosphorylated activation loop, and resolution ≤ 2.25 Å). The Ramachandran regions are marked A (alpha), B (beta), L (left), and E (epsilon). Each residue displayed in different color; X-DFG (residue before the DFGmotif, blue), DFG-Asp (green), DFG-Phe (orange) and DFG-Gly (red). (*C*) Scatterplot of side-chain dihedral angles χ_1_ and χ_2_ for DFG-Asp (green) and DFG-Phe (orange) of catalytically primed structures.

In protein kinases, the active structure and various inactive structures are distinguished by the wide variety of positions and conformations of the residues of the DFG motif. The well known DFGin and DFGout classes describe the rough position of the Asp and Phe residues of the DFG motif but fail to capture how these positions are attained. Intermediate states between DFGin and DFGout have been described (14). These three groups of structures are easily distinguished in many kinases (e.g., EGFR in Fig. 3A).

**Figure 3.**
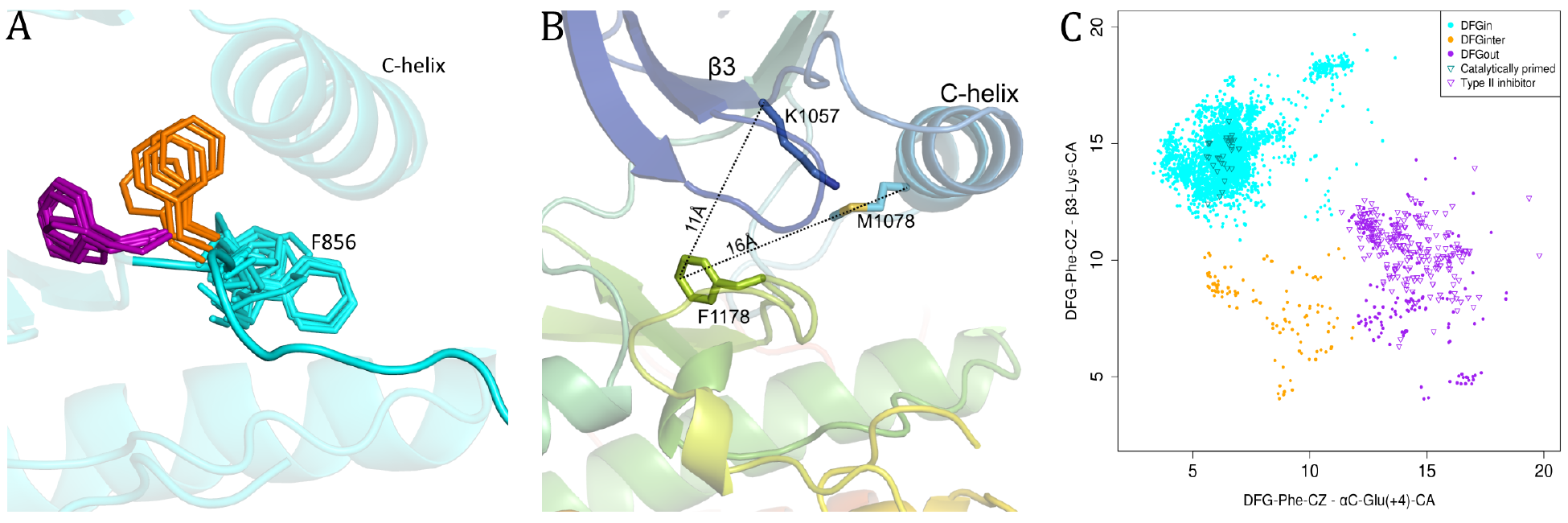
Locations of Phe side chain in DFGin, DFGinter, and DFGout structures of kinases. (*A*) Positions of DFG-Phe side chain in DFGin (cyan), DFGinter (orange), and DFGout (purple) of EGFR. (*B*) Distances, D1 and D2, used to identify the location of the Phe side chain of the DFGmotif. (*C*) Scatterplot of D1 and D2. DFGin (cyan), DFGinter (orange), and DFGout (purple) structures identified with hierarchical clustering; catalytically primed structures are marked with cyan triangles; structures with bound Type II inhibitors are marked with purple triangles.

To capture the location of DFG-Phe residue, we calculated its distance from two conserved residues in the N-terminal domain (Fig. 3B): D1 = dist(αC-Glu(+4)-Cα, DFG-Phe-Cζ); D2 = dist(β3-Lys-Cα, DFG-Phe-Cζ). D1 is the distance between the Cα atom of the fourth residue after the conserved Glu residue in the C-helix (Exxx**X**) and the outermost atom of the DFG-Phe ring (Cζ). Even as the αC helix moves outward, the hydrophobic Exxx**X** residue does not move significantly since it is located closer to the pivot point of the helix toward the back of the kinase N-terminal domain. The distance of this residue to the Phe ring serves to distinguish DFGin structures, where the Phe ring is adjacent to or under the C-helix, from DFGout structures, where the ring has moved a substantial distance laterally from the C-helix. D2 is the distance between the Cα atom of the conserved Lys residue from the β3 strand to the Cζ atom of the DFG-Phe side chain. It captures the closeness of DFG-Phe to the N-lobe β-sheet strands, thus giving an estimate of the upward position of the Phe ring, distinguishing DFGin conformations from structures where the Phe ring is in an intermediate position between DFGin and DFGout.

These distances are plotted against each other in Fig. 3C. We have clustered these distances into three groups using average linkage hierarchical clustering. The choice of three groups in clustering algorithm was guided by the visual inspection of large number of structures suggesting three broad regions or pockets occupied by DFG-Phe residue (Fig. 3A). Based on this we have classified the kinase structures into the following three groups:

a. DFGin: This is the largest group, consisting of 4,333 chains (89.6%) from 227 kinases shown in cyan-colored points in Fig. 3C, representing the DFG motif orientations where DFG-Phe is packed against or under the C-helix (cyan in Fig. 3A). It consists of many related conformations with the typical DFGin active orientation forming the largest subset of this group. All the catalytically primed structures belong to this group (cyan colored triangles in Fig. 3C).
b. DFGout: This is the second largest group, consisting of 388 chains (8%) from 60 kinases, displayed in purple-colored points representing the structures where DFG-Phe is moved into the ATP binding pocket (purple in Fig. 3A). The structures with a Type II inhibitor bound form a subset of this group (purple colored triangles in Fig. 3C).
c. DFGinter (DFGintermediate):This is the smallest group, consisting of 113 chains (2.3%) from 27 kinases, in which the DFG-Phe side chain is out of the C-helix pocket but has not moved completely to a DFGout conformation (orange in Fig. 3A, orange dots in Fig. 3C). In most of these cases DFG-Phe is pointing upwards towards the β-sheets dividing the active site into two halves. Dodson and coworkers had previously referred to this state in Aurora A kinase as ‘DFGup’ (14). Recently, Ung and coworkers have also identified this set of conformations and labeled them as ‘ωCD’ (12).

### Clustering kinase conformations based on the backbone of activation loop

For the DFG-Phe side chain to exhibit such wide-ranging localization within the kinase domain fold, the backbone and side-chain dihedral angles leading up to the Phe side chain must be divergent. By examining a large number of structures, we observed that the structural variation of the activation loop begins with the residue that precedes the DFG motif (‘X-DFG’). The position of the Phe side chain is then determined by the backbone dihedrals φ and ψ of X-DFG, DFG-Asp and DFG-Phe as well as the first side-chain dihedral of the DFG-Phe residue.

The high-resolution catalytically primed structures have very precise values of these dihedral angles (Fig. 2B and 2C). The backbone conformations of the X-D-F residues of catalytically primed structures occupy the beta (B), left (L), and alpha (A) regions of the Ramachandran map, respectively (as defined in Fig. 2B). The DFG-Phe side chain adopts a χ_1_ gauche-minus rotamer (χ_1_~-60°; χ_2_~ 90°, orange dots in the χ_1_-χ_2_ scatterplot in Fig. 2C) and points slightly downward into a pocket underneath the C-helix (Fig. 2A). In all of these structures, the DFG-Asp adopts a χ_1_ trans rotamer (~180°) and the χ_2_ dihedral is ~0° (green dots in Fig. 2C). This places the Asp carboxylate atoms in a horizontal orientation to chelate an Mg^2+^/Mn^2+^ ion on one side and to form a hydrogen bond with the NH of DFG-Gly on the other. The Mg^2+^ ion forms a tight interaction with an oxygen atom on the β-phosphate group (Fig. 2A).

Based on these observations on the catalytically primed structures, we decided to cluster the conformations of kinase structures using a metric based on the backbone dihedrals φ and ψ of X-DFG, DFG-Asp and DFG-Phe as well as the first side-chain dihedral of the DFG-Phe residue. The backbone of the DFG-Gly residue exhibits high flexibility and therefore was not included in the clustering. Each kinase chain is represented by a vector of these seven dihedrals. The distance between these vectors is calculated by a metric from directional statistics (15), which we used previously in our work on clustering antibody CDR loop conformations (16). The distance matrix calculated with this metric was used as input to DBSCAN (Density-based spatial clustering of applications with noise) which is a density-based clustering algorithm (17). DBSCAN groups together data points that are connected by high density with each other while identifying the points in low density regions as noise. We clustered the DFGin, DFGout, and DFGinter groups of structures separately.

**Figure 4.**
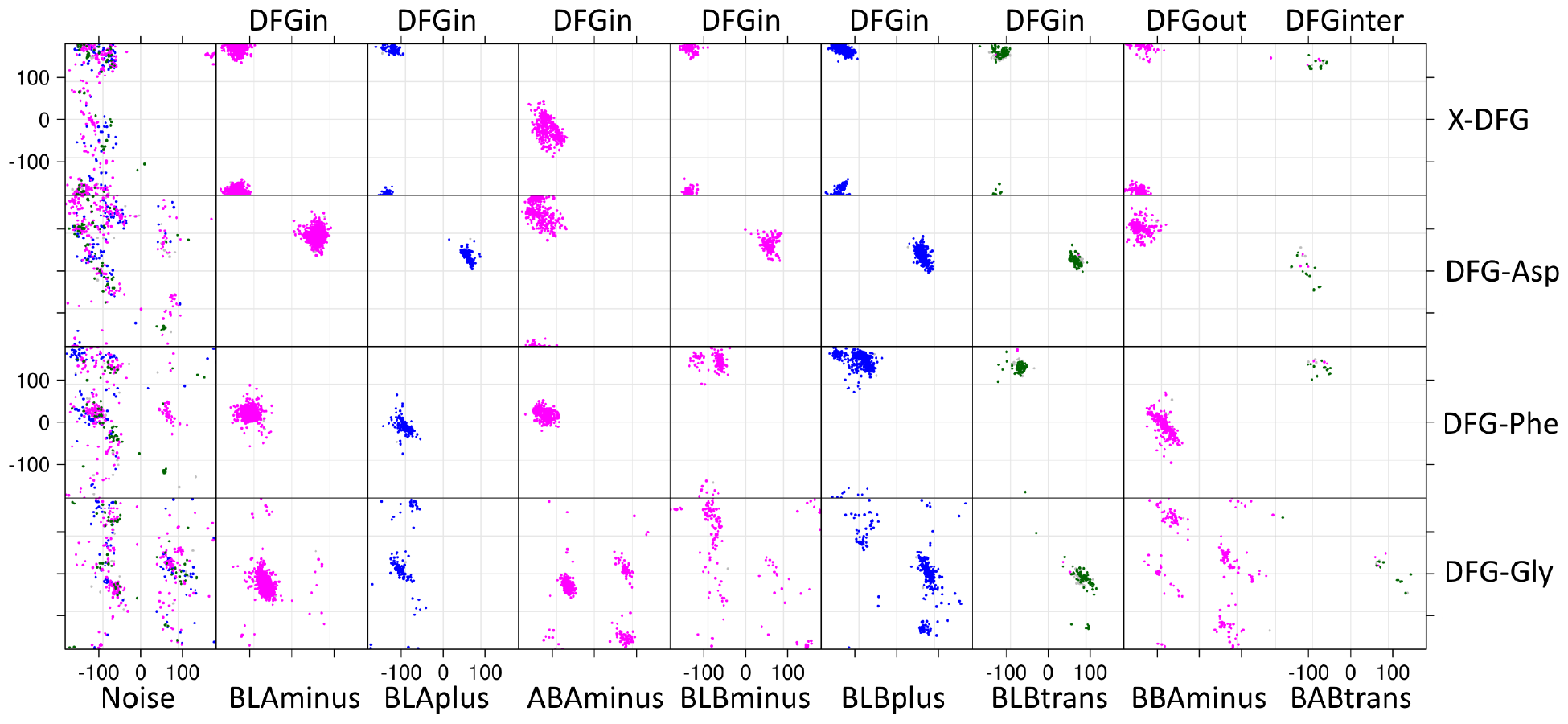
DBSCAN clustering of dihedral angles that position the DFG-Phe side chain. Clustering was performed on the DFGin, DFGinter, and DFGout groups separately with an angular metric of the backbone dihedral angles of the X-DFG, DFG-Asp, and DFG-Phe residues and the χ_1_ of DFG-Phe. Clusters are named by the Ramachandran regions of the X-DFG, DFG-Asp, and DFG-Phe residues and the χ_1_ rotamer of DFG-Phe — g^-^ rotamer (minus, magenta); g^+^ rotamer (plus, blue); trans rotamer (green).

For the DFGin group we obtained six clusters with DBSCAN (Fig. 4). The Ramachandran map dihedral angles and side-chain dihedral angles naturally cluster in high-density regions, and DBSCAN readily identified conformations of the X-DFG motif residues that conform to well known populations (18). By using the Ramachandran region annotation (A, B, L, and E) for the X, D, and F residues and the DFG-Phe χ_1_rotamer (minus = −60°; plus = +60°; trans = 180°), these clusters are labeled as BLAminus, BLAplus, ABAminus, BLBminus, BLBplus, and BLBtrans. Example structures are shown in Fig. 5A-5F. All the catalytically primed structures are observed in the BLAminus cluster; the Gly conformation for these residues is uniformly in the alpha (A) position (Fig. 2B). Although the DFG-Gly residue dihedrals were not used in the clustering, for most of the clusters there is one dominant conformation of this residue.

For the DFGout group we obtained just one cluster. In this cluster, the X-D-F residues occupy the B-B-A regions of the Ramachandran map (Fig. 4 and Fig. 5G) and DFG-Phe is in a −60° rotamer, while the Gly residue occupies all four Ramachandran conformations (A, B, L, and E). The cluster is therefore labeled BBAminus. Eighty-two percent of 244 Type II inhibitor-bound chains in 187 PDB entries of 42 different kinases are observed in this cluster; the remainder are in the DFGout noise group.

**Figure 5.**
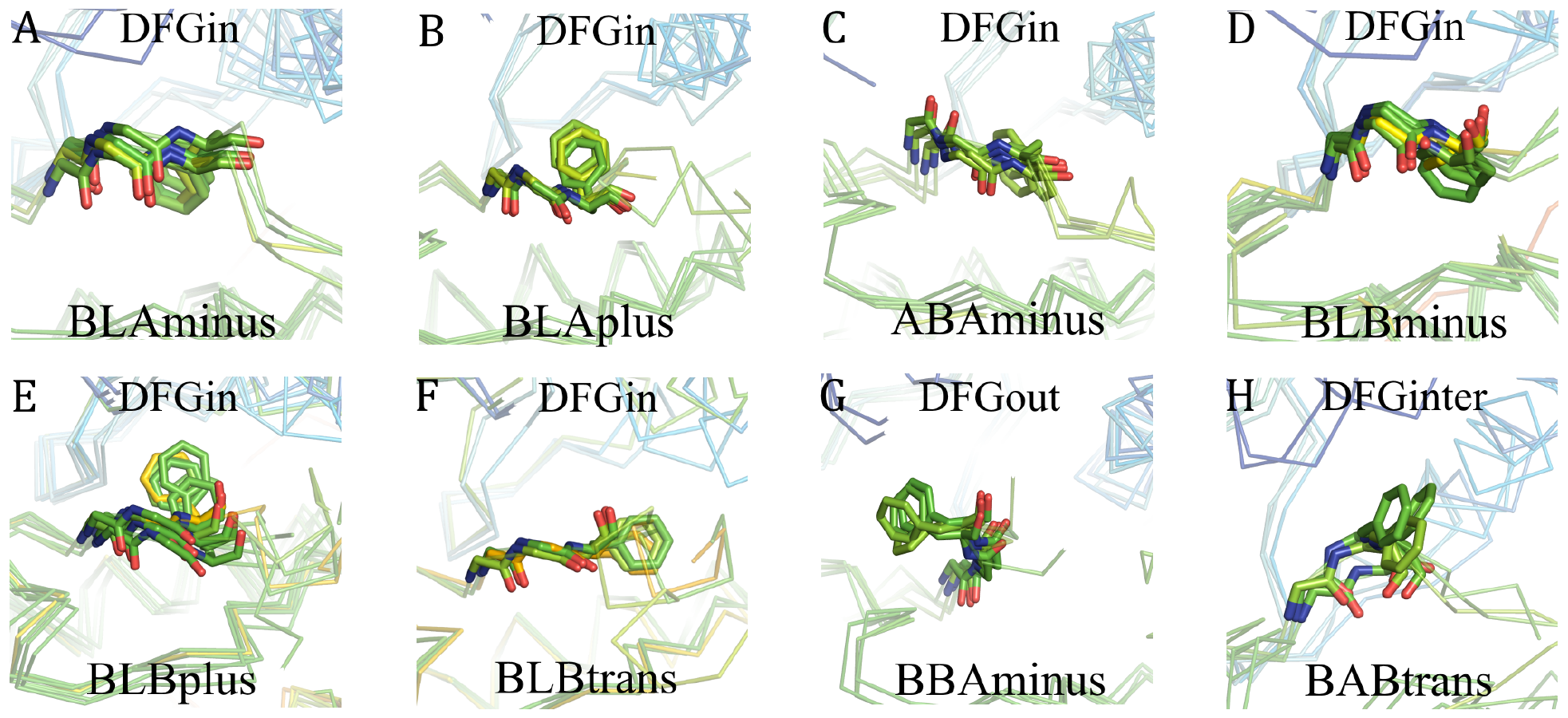
Structure examples of each cluster. (*A*) BLAminus; (*B*) BLAplus; (*C*) ABAminus; (*D*) BLBminus; (*E*) BLBplus; (*F*) BLBtrans; (*G*) BBAminus (DFGout); (*H*) BABtrans (DFGinter).

The structures in the DFGinter conformation display more variability than the other states. For the DFGinter group we obtained only one cluster of 20 chains from 8 kinases. The X-D-F residues are in a B-A-B conformation (Fig. 4 and Fig. 5H) and the DFG-Phe residue is observed in a trans rotamer with a few chains displaying a rotamer orientation between g-minus and trans (6 chains of CDK2 with DFG-Phe χ_1_ ~ −100°). The Gly residue is in an L conformation. Owing to the more prominent side-chain orientation we have labeled this cluster BABtrans.

We assigned 110 noise points to backbone clusters if the distance between these points and the nearest cluster centroid was less than a certain cutoff (Methods). The resulting clusters can be validated by their silhouette scores (Fig. S1). This leaves us with a total of 447 chains (9%) which could not be assigned to any backbone clusters. Although these structures do not get a backbone cluster label, they still belong to a specific spatial group: DFGin (48% of 447 chains), DFGout (31%), or DFGinter (21%). Cluster assignments and associated data for all human kinase chains in the PDB (except those missing the X-DFG residues or mutants thereof) are listed in Dataset S1. Mean dihedral angles and representative structures for each cluster are given in Table S2.

**Table 1:** Number of human kinases in different conformations from each phylogenetic group

| Spatial group | Clusters | All (N=4834) | Filtered (N=1621) | AGC (28) | CAMK (31) | CK1 (10) | CMGC (35) | STE (29) | TKL (18) | TYR (59) | OTHER (30) | TOTAL (244) |
| --- | --- | --- | --- | --- | --- | --- | --- | --- | --- | --- | --- | --- |
| DFGin | BLAminus | 55.6% | 58.6% | 22 | 23 | 10 | 29 | 17 | 14 | 35 | 26 | 177 |
|  | BLAplus | 3.1 | 2.4 | 2 | 1 | 0 | 1 | 3 | 1 | 7 | 2 | 17 |
|  | ABAminus | 9.1 | 7.7 | 9 | 7 | 2 | 6 | 4 | 2 | 15 | 6 | 51 |
|  | BLBminus | 3.8 | 5.0 | 0 | 3 | 0 | 4 | 4 | 3 | 8 | 4 | 26 |
|  | BLBplus | 9.5 | 9.9 | 1 | 0 | 0 | 8 | 10 | 3 | 19 | 2 | 43 |
|  | BLBtrans | 4.1 | 3.8 | 0 | 1 | 0 | 1 | 1 | 0 | 3 | 1 | 7 |
|  | Noise | 4.4 | 2.4 | 4 | 5 | 0 | 9 | 9 | 4 | 22 | 11 | 64 |
| DFGout | BBAminus | 5.1 | 5.5 | 1 | 2 | 1 | 6 | 3 | 4 | 25 | 2 | 44 |
|  | Noise | 2.9 | 2.7 | 4 | 1 | 0 | 4 | 7 | 2 | 20 | 4 | 42 |
| DFGinter | BABtrans | 0.4 | 0.2 | 0 | 0 | 0 | 0 | 1 | 0 | 6 | 1 | 8 |
|  | Noise | 1.9 | 1.3 | 2 | 7 | 0 | 1 | 3 | 0 | 6 | 2 | 21 |

Table 1 summarizes the features of each cluster. Kinase structures are most commonly observed in the BLAminus conformation (55.6% of 4,834 chains). The catalytically primed structures discussed earlier are a subset of this cluster. The next most frequent conformations are BLBplus and ABAminus observed in 9.5 and 9.1% of kinase chains respectively. However, as discussed below, some of the structures in the ABAminus state are probably incorrectly modeled. A total of 4.1% of structures have a BLBtrans orientation but a large number of structures in this cluster are only from CDK2 kinase (160 of 199 chains). The remaining DFGin clusters are BLBminus (3.8) and BLAplus (3.1%). The two largest, inactive DFGin clusters are larger than the DFGout BBAminus cluster, which represents 5.1% of kinase chains. Although there are 113 chains (2.3% of all chains) observed in the DFGinter conformation, only 20 of them cluster into the BABtrans state (0.4%).

We examined the distribution of kinase domain sequences and kinase families in our 8 clusters (domains in proteins with two kinase domains, such as the JAK kinases, are counted separately). The DFGin, DFGinter, and DFGout spatial groups are observed in 227, 27, and 68 kinase domains respectively. A total of 177 kinases with structures from all eight kinase families have been solved in the BLAminus conformation (Table 1). The other prominent DFGin clusters, ABAminus and BLBplus, were observed in 51 and 43 kinases respectively. While all the families have structures in the ABAminus cluster, BLBplus does not have structures from kinases in the CAMK or CK1 families. Forty-four kinases have structures solved in the DFGout BBAminus state; 25 of these are from the tyrosine kinase family. Among the eight subgroups, tyrosine kinase structures are the most diverse with a significant number of kinases have structures determined in all 8 conformational states. Out of 244 human kinases with known structures, 187 are solved in only one conformation.

### Validation of kinase conformational clusters with electron density

Our clusters depend on the precise directionality of the main-chain conformations of the X-D-F residues at the beginning of the activation loop of kinases. Segments of protein structures are sometimes incorrectly modeled within the electron density such that the peptide plane is flipped by 180° resulting in large dihedral angle changes of neighboring residues (19, 20). Peptide flips change ψ of residue *i* and φ of residue *i*+1 by about 180° while the other dihedrals remain approximately the same. In particular, we were concerned with some ABAminus structures that we observed to have poor electron density for the O atom of the X residue preceding the DFG motif (Fig. 6A and Fig. S2). In Fig. 6A, the negative electron density contours of the Fo-Fc maps in red show that the O atom of the X residue has been modeled without electron density support, while the positive density in green indicates the positions where an atom is missing. Both of these features point to erroneous modeling of atoms in these structures.

**Figure 6.**
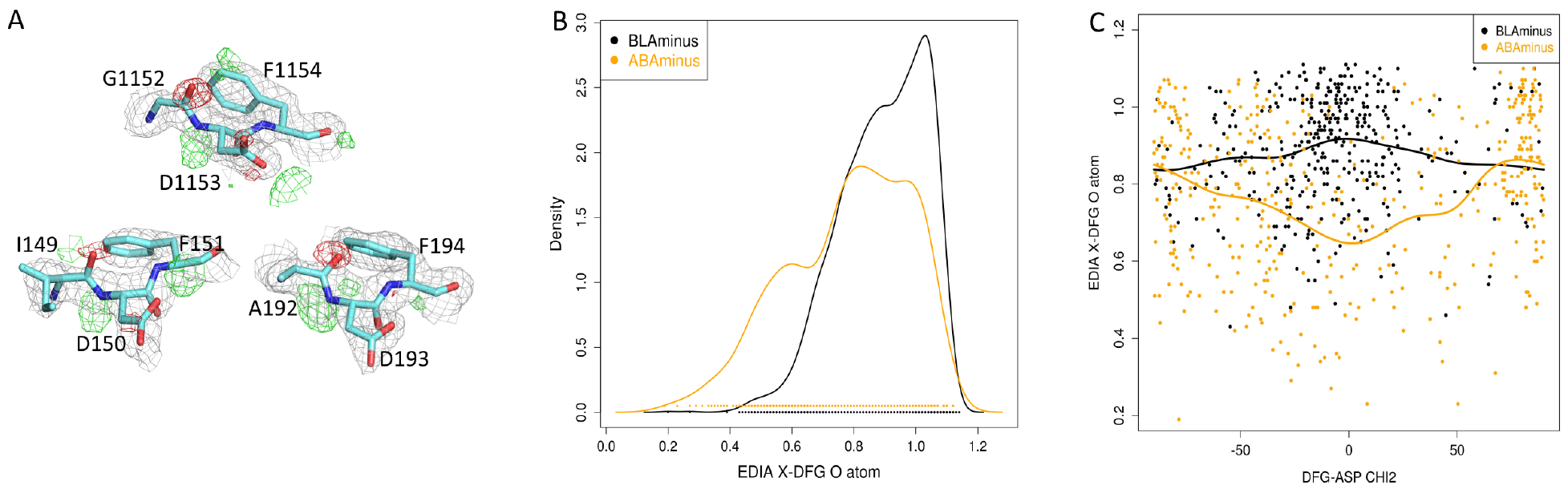
Electron densities indicate incorrectly modeled ABAminus structures. (*A*) Electron density of ABAminus structures consistent with BLAminus structure. 2Fo-Fc (gray); Fo-Fc (green=positive density, indicating density not represented by an atom; red=negative density, indicating density where an atom has been placed without electron density support. PDB ids 1K3A_A (IGF1R); 4BL1_A (MELK); 3IEC_D (MARK2). (*B*) Kernel density estimates of the EDIA scores (Electron density support of individual atoms) of backbone carbonyl of the X-DFG residue for ABAminus (orange) and BLAminus (black) structures. (*C*) Scatterplot of χ_2_ of DFG-Asp and EDIA of X_O of ABAminus (orange) and BLAminus (black) structures. BLAminus structures typically have χ_2_ near 0°. ABAminus structures with χ_2_ near 0° have poor electron density and may be mismodeled BLAminus structures. The curves are von Mises kernel regressions of the EDIA score as a function of χ_2_

We have quantified this error by using the EDIA program (‘electron density score for individual atoms’) (21). EDIA computes the weighted mean of electron density over grid points in a sphere around the atom large enough to detect both unmodeled electron density and extra modeled density where an atom is placed incorrectly. A value of 0.8 or more reflects a good electron density fit; values lower than 0.8 indicate a problem in the model. The only atoms with average EDIA scores below 0.8, consistent with a large population of mismodeled structures, are the O atoms of the X residue of ABAminus structures (Fig. 6B) and the O atom of the Phe residues of BLAplus structures (Fig. S3). This is consistent with a significant population of ABAminus structures that should have been modeled as the more common BLAminus conformation (180° change in ψ_x_ and φ_d_), and BLAplus structures that should have been modeled as the much more common BLBplus conformation (180° change in ψ_F_ and φ_g_).

A visual inspection of the electron density in mismodeled ABAminus structures suggested that the error in modeling is more commonly encountered in structures with a DFG-Asp χ_2_ dihedral of about 0°. This value of χ_2_ leads to a superposition of the Asp carboxylate in ABAminus and BLAminus structures. To examine this, for the ABAminus structures, we plotted the EDIA score of the O atom of the X residue versus χ_2_ of the Asp side chain (Fig. 6C). In correctly modeled ABAminus structures, the Asp χ_2_ dihedral is around 90°, and the EDIA scores of ABAminus and BLAminus structures are approximately the same, while in structures with poor electron density it is around 0° with EDIA scores significantly lower (min average EDIA = 0.63) than BLAminus structures with similar dihedral angles (max average EDIA = 0.92).

With these results, we have produced a filtered data set consisting of structures with resolution ≤ 2.25 Å, R-factor ≤ 0.25, and EDIA scores above 0.8 for all backbone atoms of the XDF motif (Dataset S2). All further analysis of the clusters was performed for this filtered data set. Table 1 demonstrates that the frequency of ABAminus and BLAplus in the filtered set decreases as expected, as does the noise (from 9.2% to 6.4%).

### What’s wrong with inactive structures?

The kinase active site requires a number of moving parts to be placed precisely for catalysis (Fig. 2A). We examined several structural features across our clusters to determine what is commonly missing for activity in the inactive (non-BLAminus) clusters: 1) an orientation of the C-helix such that the C-helix-Glu/β3-strand-Lys salt bridge can form; the lysine side chain must be in a specific position to form a hydrogen bond with an oxygen atom of the α phosphate and the oxygen atom bridging the β and γ phosphates of ATP; 2) the position and orientation of the Asp residue of the DFG motif; in catalytically primed structures, the Asp backbone NH positions a water molecule that forms hydrogen bonds with the adenine ring of ATP, while the carboxylate Oδ2 atom positions the Mg^2+^ ion which interacts with the β phosphate of ATP; 3) an extended activation loop is required for binding of substrate to the kinase active site; in inactive structures, the activation loop is folded up, blocking access of the active site to substrates. The results of this analysis are presented in Table 2.

**Table 2.** What’s wrong with inactives

| Cluster | Lys/Glu | AspNH | Loop | All |
| --- | --- | --- | --- | --- |
| BLAminus (951) | 94.6 | 99.7 | 99.1 | 93.9 |
| BLAplus (39) | 71.7 | 97.4 | - | - |
| ABAminus (126) | 89.6 | - | 45.2 | - |
| BLBminus (82) | 48.7 | 100 | 1.2 | 1.2 |
| BLBplus (161) | 22.3 | 100 | - | - |
| BLBtrans (62) | - | 100 | - | - |
| noise (40) | 45 | 50 | 2.5 | 2.5 |
| BBAminus (90) | 75.5 | 95.5 | 8.8 | - |
| noise (45) | 75.5 | 71.1 | 6.6 | - |
| BABtrans (4) | 25 | 75 | 25 | - |
| noise (21) | 52.3 | 61.9 | - | - |

Lys/Glu=formation of salt bridge possible (Chelix in);AspNH = DFG-Asp-NH in right position to interact with water molecule that forms hydrogen bond with ATP, as measured by percent hbond of X-DFG carbonyl with HRD-His side chain; Loop=substrate binding possible because activation loop is extended; All=percent of structures where all 3 features are present

We defined C-helix-in and C-helix-out structures (those that can or cannot form the Glu/Lys salt bridge) as those with a distance between the C-helix-Glu-Cβ and β3-Lys-C β atoms ≤ 10 Å or > 10Å respectively. In structures with an intact Glu/Lys salt bridge, 98% of the Cβ−Cβ distances are less than 10 Å (Fig. S4). This distance is therefore characteristic of the ability of the kinase to form the Glu-Lys salt bridge, regardless of whether the side-chain atom positions of these residues were correctly resolved in the structure. In the BLAminus cluster, structures in the filtered data set, which includes the catalytically primed conformation of kinases, the C-helix is in an inward orientation in 95% of the structures. Among the other DFGin clusters, ABAminus has the highest frequency of chains in a C-helix-in conformation (89%). By contrast, the large BLBplus cluster is strongly associated with a C-helix-out conformation (77%). In this cluster the g-plus rotamer of DFG-Phe points upwards, pushing the C-helix outward. In 75% of the DFGout structures in the BBAminus cluster, the C-helix remains in an inward disposition, suggesting that Type II inhibitors do not push the helix outward despite occupying a pocket adjacent to the C-helix.

The position and orientation of the Asp carboxylate enable the chelation of magnesium ion in catalytically primed structures. This conformation is stabilized by a hydrogen bond between the carboxylate Oδ1 atom and the NH of DFG-Gly. In addition, the NH atom of the Asp backbone forms a hydrogen bond with a water molecule which interacts with another water molecule which in turn forms hydrogen bonds with the adenine N7 atom (Fig. 2A). The position of the Asp backbone and side chain can be assessed in part by the presence of a hydrogen bond between the carbonyl of the X-DFG residue and the histidine side chain of the HRD motif. When this hydrogen bond is present, the NH of Asp points upwards so that it can form interactions with water molecules and the adenine ring. If the Asp side chain reaches the correct rotamer, it is able to form interactions with magnesium, which then interacts with ATP (Fig. 2A). We used the presence of a hydrogen bond between the X-DFG carbonyl atom and the HRD-His-Nɛ2 atom as a proxy for proper positioning of DFG-Asp (Table 2). All of the clusters except ABAminus have this hydrogen bond in a majority of structures. Because the carbonyl of X-DFG points upwards in ABAminus, the NH of DFG-Asp points downwards and is unable to position water molecules that interact with the adenine moiety of ATP. As we observed in Fig. 6C, the aspartic acid side chain has a χ_2_ dihedral angle of near 90° in most ABAminus structures with good electron density. In this orientation, DFG-Asp would not bind magnesium ion without a rotation of χ_2_ which is unfavorable in this combination of backbone conformation and χ_1_ rotamer (22).

An extended conformation of the activation loop is required for substrate binding to kinases, since the more folded loop structures block the substrate binding site. When the activation loop is extended, the backbone N atom of the sixth residue in the loop (DFGxxX) makes a hydrogen bond with the backbone O atom of the residue preceding the HRD motif (X-HRD, Fig. 2A). Using this criterion, our analysis shows that 99% of chains in the BLAminus cluster have an extended activation loop in the filtered data set (Table 2). Among the other DFGin clusters, 45% of ABAminus chains have their activation loop in a similar conformation. Beyond BLAminus and ABAminus, an extended activation loop is rare.

If we take these three criteria into account, only the BLAminus structures are capable of binding ATP, magnesium ion, and substrate simultaneously with 94% of structures passing all three (Table 2). The only structures from other clusters that pass the three criteria consist of one DFGin noise structure and one BLBminus structure, both from the same PDB entry, human IRAK4, PDB entry 2NRY, chains A and B respectively. Notably, the other two monomers in this crystal are BLAminus structures. Those chains also exhibit poor electron density for the residues after DFG.

To add some additional detail to the structural features of our clusters, we examined them for the side-chain conformation of the DFG-Asp side chain and the presence of beta turns (Table S3), and the position of the Gly-rich loop (Fig. S5). The DFG-Asp side chain has a predominant conformation in each of our clusters. Ninety-six percent of BLAminus structures possess a Type I beta turn beginning with DFG-Phe (sequence FGXX) with a hydrogen bond between the carbonyl of DFG-Phe and the NH of the DFG+2 residue. This beta turn determines the path of the activation loop into an extended conformation suitable for substrate binding. Fifty-five percent of ABAminus structures contain this beta turn but no other clusters do. The other clusters contain a beta turn beginning at DFG-Asp, including 60% of BLBplus structures that have a Type II turn (sequence DFGX) and 49% of BLAplus structures with a Type I turn at this position.

### Conformational properties of kinases with specific ligands

The distribution of ATP-bound and apo structures across our clusters may give an indication of what inactive conformations some kinases prefer in vivo, although we are limited by the structures determined for each kinase and the conditions employed during crystallization. Table 3 lists the percent of ATP-bound (or analogue), inhibitor-bound, or apo structures in each of our clusters for the filtered data set. Seventy-six percent of ATP-bound structures are in the BLAminus cluster. A triphosphate is never found in complex with a kinase in the DFGout and DFGinter states because the location of the DFG-Phe side chain would make the binding of ATP unfavorable. Among ATP+Mg bound structures, only the BLAminus, BLBplus, BLBtrans, and DFGin-noise states are represented. As expected, there are no ABAminus structures with bound magnesium in the filtered data set.

**Table 3.** Distribution across clusters for different types of ligands

| Cluster | Apo<br>(N=149) | ATP<br>(118) | ATP+Mg<br>(60) | Inhi.<br>(1308) |
| --- | --- | --- | --- | --- |
| BLAminus | 64.4 | 76.2 | 68.3 | 58.7 |
| BLAplus | 4 | - | - | 2.5 |
| ABAMinus | 6.7 | 1.6 | - | 8.8 |
| BLBminus | 4 | - | - | 5.8 |
| BLBplus | 5.3 | 5.9 | 11.6 | 11.3 |
| BLBtrans | 2.6 | 11.8 | 20 | 3.4 |
| noise | 2.6 | 3.3 | - | 2.4 |
| BBAMinus | - | - | - | 6.9 |
| noise | 5.3 | - | - | 2.8 |
| BABtrans | 1.3 | - | - | - |
| noise | 3.3 | - | - | 1.2 |
| Total | 100% | 100% | 100% | 100% |

ATP=Triphosphates (ATP, ANP, ACP and AGS); ATP+Mg=both Triphosphate and Mg bound. “-” = none.

The apo-form kinases are predominantly in the BLAminus state (64%); the other DFGin structures comprise 25% of the apo structures. This shows that multiple clusters which are observed within the DFGin group are likely to be naturally occurring states even in the absence of any bound inhibitor. However, relatively few apo structures of kinases exist in the DFGout cluster (BBAminus, 0%) or the DFGinter cluster (BABtrans, 1.3%) and as noted above ATP-bound structures do not either. These distributions are in contrast to inhibitor-bound structures, which are observed across all the conformations. A total of 239 chains (96%) in the BBAminus cluster are in complex with an inhibitor, 200 chains (80%) of which are in complex with a Type II inhibitor (listed in Dataset S3).

**Figure 7.**
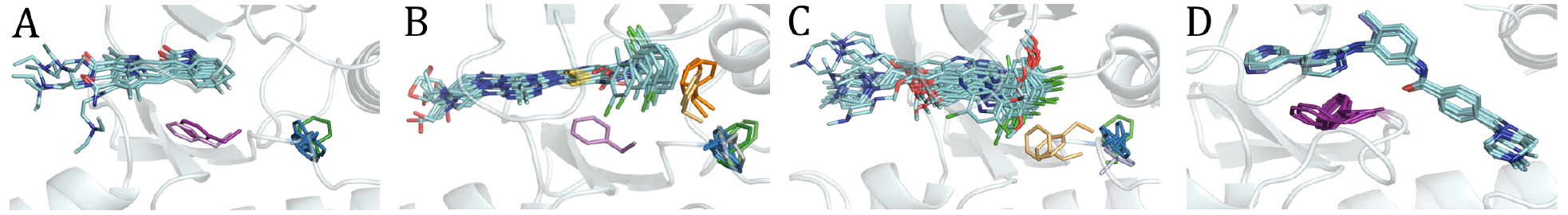
Inhibitors bound to multiple conformational clusters. Color code for different states is BLAminus (sky blue), BLAplus (dark green), BLBminus (light gray), BLBplus (light green), BBAminus (purple), BABtrans (orange), DFGout-noise (light purple), and DFGinter-noise (light orange). (A) Sunitinib bound to PAK6, PHKG2, and STK24 in BLAminus; ITK in BLAplus; CDK2 in BLBminus; KIT in BBAminus; KIT in DFGout-noise. (B) Dasatinib bound to ABL1, STK10, and STK24 in BLAminus; BTK in BLAplus; EPHA2 in BLBminus; PTK6 in BLBplus; BMX and BTK in BABtrans; ABL1 in DFGinter-noise; DDR1 in DFGout-noise. (C) Bosutinib bound to EPHA2, PMYT1, SRC in BLAminus; ERBB3 in BLBplus; STK10 in DFGin-noise; ABL1 and KCC2A in DFGinter-noise. (D) Imatinib bound to ABL1, ABL2, CSF1R, DDR1, KIT, and LCK in BBAminus.

Our clustering scheme makes the labeling of kinase conformations straightforward (Fig. 7). For example, sunitinib is bound to several DFGin conformations: STK24, PAK6, and PHKG2 in BLAminus conformations; ITK in a BLAplus conformation; and CDK2 in a BLBminus conformation (Fig. 7A). It is also bound to tyrosine kinases KIT and VEGFR2 in DFGout-BBAminus conformations. Dasatinib binds to different conformations of the same kinase in different PDB entries: ABL1 in BLAminus and BABtrans states, and BTK in BLAplus and BABtrans conformations. It also binds to BLAminus conformations of ABL2, PMYT1 SRC, EPHA4, STK10, and STK24 as well as BLBplus in PTK6, BLBminus in EPHA2, and DFGinter-BABtrans structures in BMX (Fig. 7B). Fig. 7C shows similar results for bosutinib. As a type II inhibitor, imatinib binds to DFGout structures in the BBAminus state (Fig. 7D). This kind of analysis makes clear that some inhibitors do not bind to all kinases in the same conformational state or even to one kinase in only one conformational state. It also argues that classifying inhibitors by the state of the kinase they bind to is not necessarily useful. It may be more productive to classify them by what volumes within the kinase active and C-helix sites they occupy (8).

### Comparison with previous kinase classification schemes

We compared our labels to three previously published classification schemes. Taylor and coworkers have defined a regulatory spine as a stacking of four hydrophobic residues which dynamically assemble in the active state of the kinase (23). These consist of the HRD-His, the DFG Phe, C-helix Glu+4, and a residue in the loop just prior to strand β4 (Fig. 8). Taylor and coworkers defined the regulatory spine only by comparison of structures; they did not define what constitutes the presence or absence of the spine in any one structure. We took a simple approach, and we define the regulatory spine as present if the minimum side-chain/side-chain contact distances among these four residues (1–2, 2–3, and 3–4) are all less than 4.5 Å. The regulatory spine is present in 98% of the BLAminus structures and 100% of the BLAplus structures, but it is also present in about 70% of BLAplus, ABAminus, BLBminus, and BLBplus clusters, indicating that its presence is not sufficient for defining active kinase structures (Fig. 8 and Table S4). Rather it is a feature of most DFGin structures, whether they are active or inactive enzymes. The regulatory spine is never intact in DFGout and DFGinter structures, because the DFG-Phe residue has moved out from the back pocket, making the first distance greater than 4.5 Å in all of these structures.

**Figure 8.**
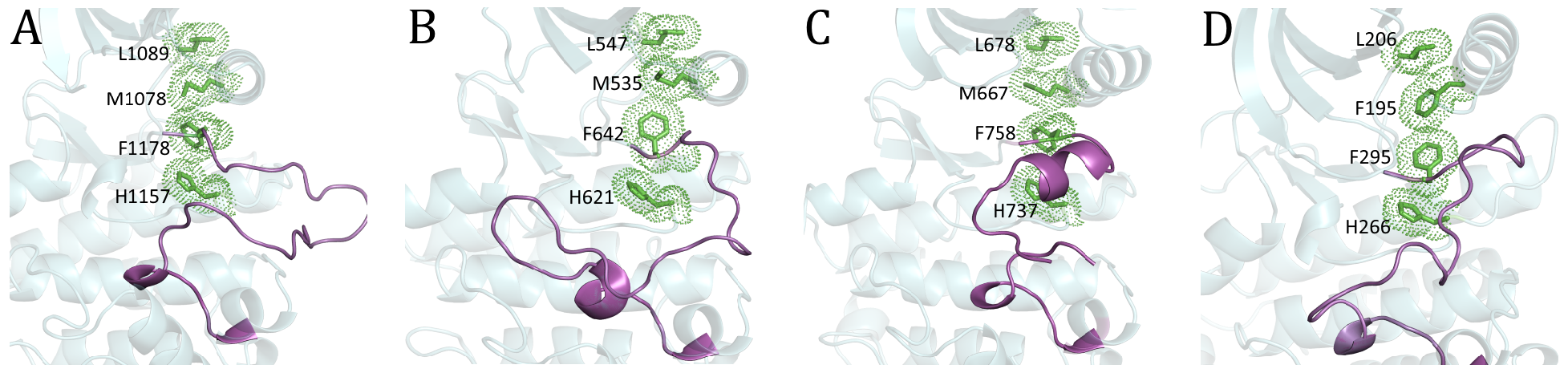
The intact regulatory spine of active and inactive structures. (A) active BLAminus (3BU5_A, INSR); (B) inactive BLAplus (4F64_A, FGFR1); (C) inactive BLBminus (5NK7_A, EPHA2); D) inactive BLBplus (4UY9_B, M3K9). We consider the spine intact if there is at least one atom-atom contact between the side chains of each residue of the spine within 4.5 Å.

Möbitz published a classification scheme for kinase structures based on the position of the C-helix and two pseudo torsions of the Cα atoms of the XDFG and DFGX residues, capturing the relative orientation of DFG-Asp and DFG-Phe (11). The scheme uses DFGin and DFGout as the major labels and subdivides them into 12 conformations (Table S5). Although this scheme is able to identify most of the BLAminus structures as active, it fails to distinguish them from ABAminus. It divides DFGout structures into five groups, three of which are unintuitively named for some representative members of each cluster (‘AuP BRAF’, etc.). Further, it merges our DFGinter structures with our DFGout noise structures into a ‘DFG Flipped’ category, even though the position of the DFG-Phe residues in these two conformations are very far apart.

More recently, Ung and coworkers used two vectors to capture the orientation of the DFGmotif (DI and DO for DFGin and DFGout) and one distance to identify the inward and outward orientation of the C-helix (CI and CO), dividing kinases into four groups as CI-DI, CI-DO, CO-DI, CODO (12) (Table S6). Ung’s method successfully distinguishes DFGinter conformations, which they call ωCD. However, it fails to distinguish active from inactive DFGin structures, and lumps our six DFGin clusters into the CI-DI and CO-DI clusters based purely on the position of the C-helix. But as our clustering has shown, these conformations have different backbone and DFG-Phe side-chain orientations and positions (Fig. 5A-5F).

## DISCUSSION

We have developed a clustering and labeling scheme which first divides the kinase structures into three groups, based on the location of the DFG-Phe side chain, that are further clustered based on the orientation of the activation loop. To cluster the orientation of the activation loop, we have used the dihedral angles that determine the placement of the Phe side chain: the backbone dihedrals (φ, ψ) of the X-D-F residues and the first side-chain dihedral (χ_1_) of the DFG-Phe residue. These are parameters used to define the conformation of any polypeptide chain. From this clustering, we have developed a simple nomenclature for kinase conformations that is intuitive and easily applied by structural biologists when they determine a new kinase structure. It is based on the region occupied by the backbone dihedrals on the Ramachandran plot and the side-chain rotamer of DFG-Phe.

One of the most important results of our clustering is that it is able to identify several distinct states within the ensemble of active and inactive DFGin structures, which have usually been grouped together in previous clustering schemes (9, 10, 12). We have determined that the most frequently observed conformation, BLAminus, is also the active state conformation of kinases. Catalytically primed structures, those containing bound ATP and Mg^2+^/Mn^2+^ ion and a phosphorylated activation loop, are all members of the BLAminus cluster. We find that nearly all BLAminus structures have structural features consistent with an active kinase.

Among the inactive states in the DFGin group, BLBplus and ABAminus are the most frequent conformations with almost the same frequency at 9.5 and 9.1% respectively. However, we observed that many structures with ABAminus conformations are likely to be incorrectly modeled. In these structures, the peptide group spanning the X and D residues of the X-DFG sequence is flipped such that the backbone carbonyl oxygen of the X-DFG residue is misplaced. This kind of error in structure determination is fairly common, in this case leading to BLAminus structures being incorrectly modeled as far less common ABAminus structures. Upon removing low-resolution and poorly determined structures, BLBplus becomes even more prevalent than ABAminus (10% and 7.7% respectively) and is the most frequently occurring inactive conformation of kinases. In this conformation, the DFG-Phe ring is underneath the C-helix but pointing upwards, and the C-helix is pushed outwards creating extra volume, a region which is sometimes exploited for inhibitor design. BLBplus is sometimes referred to as the ‘SRC-like inactive’ state (24, 25), although the latter has not been explicitly defined.

We have also examined why each type of inactive state is inactive. In the three BLB states (BLBplus, BLBminus, BLBtrans), the C-helix is pushed outwards in more than 50% of cases such that the Glu/Lys salt bridge in the N-terminal domain cannot form. In the ABAminus and DFGout and DFGinter states, the Asp side chain is not positioned to bind Mg so that it can interact with ATP. In all of the active states except ABAminus, the activation loop is not extended in a way that allows substrate binding.

We have compared our clustering and labeling scheme with three previously published methods. The regulatory spine defined by Kornev and coworkers is a commonly used method to distinguish between active and inactive states (23), although it has not been explicitly defined. Our data indicate that the presence of the regulatory spine can only reliably distinguish DFGin structures from DFGout and DFGinter structures. It fails to identify the different kinds of inactive states within the DFGin group, most of which have an intact regulatory spine. Möbitz developed a classification scheme based on the C-helix position and the XDFG and DFGX Cα pseudo dihedrals of the activation loop (11). His labeling scheme is rather complicated and the names are unintuitive and difficult to rationalize. Ung and colleagues divided kinase structures into DFGin (DI), DFGout (DO), and DFGinter (ωCD) categories and then divided the DFGin ensemble into just two states, DFGin-C-helix-in and DFGin-C-helix-out (12). This scheme characterizes the C-helix and DFG-Phe positions but it does not capture the variability of the DFGin states and fails to separate active from inactive kinase structures.

Inhibitors have been classified into Type I and Type II inhibitors, depending on whether they bind to the ATP binding site alone or to the ATP binding site and the volume next to the C helix that is exposed in DFGout structures. Our classification scheme allows us to determine that some inhibitors bind to different conformational states in different kinases, and sometimes to different states of the same kinase. In some cases, the inhibitor makes quite different contacts when bound to different states of the kinase, since the X-DFG residues may come in contact with the inhibitor. Some Type I inhibitors even bind to DFGout conformations in our BBAminus cluster. We provide a list of structures of FDA approved kinase inhibitors bound to kinases with cluster labels provided in Table S7.

Our clustering and nomenclature can be applied to interpret the dynamical properties of various conformational states of kinases and the transitions between them. For instance, Tong et al. studies SRC kinase with variants of dasatinib that stabilized three states of the kinase (26). In our nomenclature, these three states were BLAminus, BBAminus, and BLBplus. The BLBplus structure had significantly reduced dynamics of the HRD loop, the activation loop, and the loop between the F and G helices. Multiple studies have used molecular dynamics (MD) simulations to study the transition from active to inactive states in protein kinases (27–29). However, due to lack of a consistent nomenclature divergent DFGin states have been labeled as Src-like inactive. Levinson et al performed simulations of ABL1 from a ‘SRC-like inactive state’ (PDB: 2G1T) that we label a BLBplus structure. However, Shan and colleagues started simulations of EGFR from a ‘SRC-like inactive’ structure (PDB: 2GS7), which is actually BLBtrans, an infrequently observed conformation for most kinases. Our scheme for assigning structures to different conformational states will improve the analysis of molecular dynamics simulations of kinases described in these studies.

Finally, significant effort has been expended to produce comparative models of kinases in different conformational states and to study the docking of inhibitors to these structures (30, 31). A more reliable classification of the states of kinases will have a positive impact on choosing templates for producing models of kinases in various biologically and therapeutically relevant states.

## METHODS

The structures having kinase domains were identified from the file *pdbaa* (December 1, 2017) in the PISCES server (32) with 3 rounds of PSI-BLAST (33) with the sequence of human Aurora A kinase (residues 125-391) as query. This profile was used again to search *pdbaa* with an E-value cutoff of 1.0×10^-15^ to eliminate structures of proteins that are not kinases or contain divergent folds. The structures with resolution worse than 4 Å and those with missing or mutated residues in the DFGmotif were removed. The conserved motifs were identified from pairwise sequence and structure alignments with Aurora A. Clustering was performed on all chains containing a kinase domain in these entries. We have defined a filtered dataset of high-resolution protein kinase structures. This set includes structures which satisfy the following criteria: 1) Resolution better than or equal to 2.25 Å, R-factor ≤ 0.25, and no pseudokinases; 2) EDIAscore of X-DFG, DFG-Asp and DFG-Phe backbone atoms ≥ 0.8; 3) Overall EDIAscore of DFG-Asp and DFG-Phe residues (including side chains) ≥ 0.8. The pseudokinases we removed from the data set comprise CSKP, ERBB3, ILK, KSR2, MKL, STK40, STRAA, TRIB1, VRK3, WNK1, and WNK3. The filtered data set is provided in Dataset S2.

To capture the location of the DFG-Phe residue, we calculated its distance from two conserved residues in the binding site: 1) αC-Glu(+4)-Cα to DFG-Phe-Cζ; 2) β3-Lys-Cα to DFG-Phe-Cζ. We clustered structures into three groups using the sum of squares of these distances and average linkage hierarchical clustering algorithm using hclust function in the statistical software R (34).

Dihedral angle clustering of the DFGin, DFGinter, and DFGout groups was performed with DBSCAN from the fpc package in the R program (35). For each pair of structures, *i* and *j*, the distance between them was calculated as the sum of an angular distance function over 7 dihedral angles:

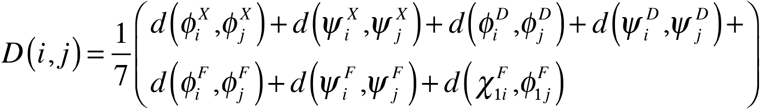

The distance for each angle is equal to the square of the chord length between the ends of two vectors originating at the center of a unit circle (15, 16): *d*(*θ*_1_,*θ*_2_) = 2(1 - cos (*θ*_2_–*θ*_1_)).

DBSCAN requires two parameters, ɛ and MinPts. Data points with at least MinPts points within a distance ɛ are considered ‘core points’. Points within ɛ of a core point but not themselves core points are called border points. All other points are considered noise. If we make a graph by treating the core points as nodes and place edges between them if they are within ɛ of each other, then the clusters are identified as the connected subgraphs of the whole graph. Border points are then assigned to the cluster that contains the closest core point to the border point. The noise points are not assigned to clusters.

The choice of appropriate parameters is critical for determining the clusters. The value of MinPts is approximately equal to the smallest cluster that the procedure will return. If ɛ is too small, some clusters will be inappropriately subdivided into many small, dense clusters. If ɛ is too large, adjacent clusters may be merged. We have an advantage that protein φ, ψ dihedral angle pairs naturally cluster within the Ramachandran map. In addition, outliers are easily identified in forbidden regions of φ, ψ space. We used these features to scan through pairs of ɛ and MinPts values for each spatial group separately, minimizing the number of points identified as noise without visibly merging clusters representing distinct basins of the Ramachandran map. We used ɛ and MinPts values of 0.05 and 20 for DFGin, 0.06 and 20 for DFGout, 0.3 and 15 for DFGinter. We validated our clusters with the silhouette metric (Fig. S1).

To assign labels to as many data points as possible, the noise points whose distance from their nearest cluster centroid is less or equal to 0.3 units (equivalent to an average dihedral angle difference of 21°) were assigned to those respective clusters. The remaining noise points were still labeled with one of the three spatial group labels.

Hydrogen bond analysis was performed with HBPlus (36). The classification of beta turns into different turn types was done using a Python program by Maxim Shapovalov (https://github.com/sh-maxim/BetaTurn18) (37). The graphs were made using various plotting functions in the statistical package R. The molecular images were created using Pymol https://www.pymol.org/). Electron densities were calculated with the program Phenix (38). Electron densities were validated with the program EDIA (21).

We have created a simple tool to assign our kinase structures in PDB format to our clusters (http://dunbrack.fccc.edu/kinasetool). A web database andserver will be forthcoming.

## ACKNOWLEDGMENTS

VM thanks Fox Chase Cancer Center for Elizabeth Knight Patterson postdoctoral fellowship. We thank Maxim Shapovalov for providing a program to identify beta turn types. This work was funded by NIH grants R01 GM084453 (R.L.D.) and R35 GM122517 (R.L.D).

---

**Table S1:** List of catalytically primed structures

| Uniprot | PDB | Chain | Reso | X_Phi | X_Psi | Asp_Phi | Asp_Psi | Asp_Chi1 | Asp_Chi2 | Phe_Phi | Phe_Psi | Phe_Chi1 | Ligand |
| --- | --- | --- | --- | --- | --- | --- | --- | --- | --- | --- | --- | --- | --- |
| AKT2 | 1O6K | A | 1.7 | -120.8 | -176.0 | 56.8 | 82.9 | 186.7 | 12.6 | -98.9 | 24.5 | 299.4 | ANP |
| AKT2 | 1O6L | A | 1.6 | -121.5 | -178.2 | 58.4 | 83.3 | 187.4 | 7.0 | -98.8 | 24.7 | 297.6 | ANP |
| AURKA | 5DN3 | A | 2.0 | -130.9 | -174.4 | 55.3 | 74.8 | 189.2 | 0.9 | -89.0 | 14.2 | 275.5 | ATP |
| AURKA | 5DNR | A | 1.9 | -132.2 | -179.7 | 60.8 | 76.7 | 188.4 | 2.7 | -90.1 | 18.7 | 275.9 | ATP |
| CDK2 | 1QMZ | A | 2.2 | -146.3 | -172.6 | 47.5 | 78.4 | 192.7 | -2.0 | -93.9 | 24.3 | 280.2 | ATP |
| CDK2 | 1QMZ | C | 2.2 | -147.5 | -169.9 | 41.1 | 80.5 | 192.6 | -4.4 | -95.5 | 25.6 | 283.1 | ATP |
| CDK2 | 4EOJ | A | 1.6 | -134.1 | 177.6 | 53.9 | 78.1 | 191.5 | 9.6 | -94.3 | 17.8 | 277.9 | ATP |
| CDK2 | 4EOM | A | 2.1 | -135.9 | -180.0 | 56.8 | 76.4 | 194.3 | -4.5 | -93.7 | 18.3 | 281.7 | ATP |
| CDK2 | 4EOM | C | 2.1 | -134.3 | 179.5 | 56.9 | 73.6 | 194.6 | 8.9 | -89.3 | 27.1 | 288.3 | ATP |
| CDK2 | 4EOO | A | 2.1 | -137.3 | -178.3 | 51.9 | 74.5 | 189.5 | -2.6 | -88.8 | 23.7 | 283.6 | ATP |
| CDK2 | 4EOO | C | 2.1 | -132.9 | -176.4 | 50.3 | 74.1 | 194.7 | 1.1 | -84.3 | 27.4 | 283.8 | ATP |
| CDK2 | 4EOQ | A | 2.1 | -136.2 | -179.0 | 55.1 | 77.3 | 197.2 | 1.6 | -91.4 | 19.3 | 282.3 | ATP |
| FGFR2 | 2PVF | A | 1.8 | -138.9 | -175.2 | 54.7 | 80.5 | 190.7 | -12.4 | -90.1 | 24.3 | 291.1 | ACP |
| FGFR2 | 3CLY | A | 2.0 | -138.8 | -172.7 | 51.3 | 79.8 | 187.4 | -3.1 | -91.3 | 32.5 | 290.5 | ACP |
| INSR | 1IR3 | A | 1.9 | -133.7 | -168.2 | 47.5 | 85.0 | 199.9 | -12.6 | -100.3 | 24.6 | 303.4 | ANP |
| INSR | 3BU5 | A | 2.1 | -128.7 | -166.6 | 44.5 | 89.6 | 189.5 | -14.9 | -102.8 | 23.9 | 295.9 | ATP |
| KAPCA | 4WB5 | A | 1.6 | -123.1 | 176.8 | 69.0 | 80.0 | 197.4 | -7.7 | -91.5 | 24.9 | 270.2 | ATP |
| KAPCA | 4WB6 | A | 2.1 | -118.4 | -169.5 | 51.7 | 82.5 | 197.0 | -10.7 | -95.6 | 17.7 | 271.2 | ATP |
| KAPCA | 4WB6 | B | 2.1 | -117.8 | -168.3 | 53.2 | 78.4 | 192.7 | -12.2 | -91.3 | 29.4 | 275.0 | ATP |
| KAPCA | 4WB8 | A | 1.5 | -121.5 | -177.2 | 62.7 | 77.8 | 196.8 | -6.9 | -92.0 | 24.6 | 268.9 | ATP |
| KPCI | 5LI1 | A | 2.0 | -129.2 | 177.0 | 73.2 | 90.2 | 187.2 | 4.7 | -103.9 | 11.5 | 308.2 | ANP |
| MK01 | 5V60 | A | 2.1 | -124.3 | -175.3 | 60.5 | 75.4 | 193.8 | 5.9 | -91.1 | 25.4 | 286.1 | ACP |
| PAK1 | 3Q53 | A | 2.0 | -131.4 | -173.5 | 59.2 | 73.6 | 202.0 | -16.8 | -91.7 | 13.0 | 288.7 | ATP |
| PAK4 | 4JDI | A | 1.8 | -134.3 | 177.9 | 63.7 | 71.6 | 185.9 | 8.0 | -94.6 | 13.8 | 289.8 | ANP |
| PDPK1 | 4A07 | A | 1.8 | -135.4 | -178.9 | 76.0 | 72.4 | 205.6 | 22.7 | -109.0 | 8.7 | 286.3 | ATP |
| PDPK1 | 4AW0 | A | 1.4 | -134.8 | 177.3 | 78.7 | 76.2 | 199.3 | 18.1 | -106.3 | 8.6 | 283.5 | ATP |
| PDPK1 | 4AW1 | A | 1.6 | -137.2 | -178.3 | 69.0 | 84.7 | 188.0 | 9.9 | -105.4 | 17.0 | 286.1 | ATP |
| STK24 | 4QML | A | 1.8 | -130.4 | 172.3 | 70.8 | 79.3 | 196.8 | -0.6 | -104.6 | 20.0 | 285.9 | ANP |

**Table S2:** Cluster centroids

| Spatial groups | Clusters | X-DFG<br>$\Phi, \psi$ | DFG-Asp<br>$\Phi, \psi$ | DFG-Phe<br>$\Phi, \psi$ | <sup>†</sup>DFG-Phe<br>$\chi_1$ | Examples |
| --- | --- | --- | --- | --- | --- | --- |
| DFG<sub>in</sub> | BLA<sub>minus</sub> | -129, 179 | 61, 81 | -97, 20 | -71 | 3BU5_A (INSR) |
|  | BLA<sub>plus</sub> | -119, 168 | 59, 34 | -89, -8 | 56 | 4OTH_A (PKN1) |
|  | ABA<sub>minus</sub> | -112, -8 | -141, 148 | -128, 23 | -64 | 1U46_A (ACK1) |
|  | BLB<sub>minus</sub> | -135, 175 | 60, 65 | -79, 145 | -73 | 1R3C_A (MK14) |
|  | BLB<sub>plus</sub> | -125, 172 | 60, 33 | -85, 145 | 49 | 3PIX_A (BTK) |
|  | BLB<sub>trans</sub> | -106, 157 | 69, 21 | -62, 134 | -145 | 3VRZ_A (HCK) |
|  | Noise | - | - | - | - | 3BZ3_A (FAK1) |
| DFG<sub>out</sub> | BBA<sub>minus</sub> | -138, -176 | -144, 104 | -82, -9 | -70 | 3BE2_A (VGFR2) |
|  | Noise | - | - | - | - | 3LW0_A (IGF1R) |
| DFG<sub>inter</sub> | BAB<sub>trans</sub> | -80, 128 | -117, 24 | -85, 133 | -179 | 3SXR_A (BMX) |
|  | Noise | - | - | - | - | 5EW9_A (AURKA) |

<sup>†</sup>Phe side chain rotamers: plus: $0^\circ < \chi_1 \leq 120^\circ$ ; trans: $\chi_1 > 120^\circ$ or $\chi_1 \leq 240^\circ$ ; minus: $240^\circ > \chi_1 \leq 360^\circ$

**Table S3:** Beta turns in activation loop across different conformations (% chains) and DFG-Asp X1 rotamer populations of each cluster (% chains)

| Spatial groups | Clusters | Beta turn beginning with DFG-Asp (turn type) | Beta turn beginning with DFG-Phe (turn type) | Plus | Trans | Minus |
| --- | --- | --- | --- | --- | --- | --- |
| DFGin | BLAminus | - | 96 (I) | - | 85 | 15 |
|  | BLAplus | 49 (I) | 11 (II) | - | 25 | 75 |
|  | ABAminus | - | 55 (I) | 27 | 71 | 2 |
|  | BLBminus | 7 (II) | - | - | 67 | 31 |
|  | BLBplus | 60 (II) | 11 (II') | - | 25 | 75 |
|  | BLBtrans | 95 (II) | - |  | 5 | 95 |
| DFGout | BBAminus | 7 (I) | 10 (II') | 31 | 66 | 3 |
| DFGinter | BABtrans | 95 (II) | - | 52 | 39 | 10 |

Aspartic acid $\chi_1$ rotamers are given as Plus ( $0 \leq \chi_1 < 120^\circ$ ), Trans ( $120^\circ \leq \chi_1 < 240^\circ$ ), Minus ( $240^\circ \leq \chi_1 < 360^\circ$ ).

**Table S4:** Structures with intact regulatory spine across different clusters (% chains)

| Spatial group | Clusters | <sup>†</sup>S1<br>(HRD-His /<br>DFG-Phe) | S2<br>(DFG-Phe /<br>αC-Glu+4) | *S3<br>(αC-Glu+4 /<br>β4-X) | All three |
| --- | --- | --- | --- | --- | --- |
| DFGin | BLAminus | 100 | 99 | 99 | 98 |
|  | BLAplus | 100 | 100 | 100 | 100 |
|  | ABAminus | 67 | 98 | 98 | 67 |
|  | BLBminus | 94 | 78 | 85 | 70 |
|  | BLBplus | 99 | 74 | 94 | 73 |
|  | BLBtrans | 99 | 27 | 95 | 27 |
| DFGout | BBAminus | - | - | 93 | - |
| DFGinter | BABtrans | - | 100 | 100 | - |

<sup>†</sup>A contact is defined if the minimum distance between the side-chain atoms of the two residues is less than or equal to 4.5Å.
\* β4-X is typically a hydrophobic residue from β4 constituting the regulatory spine (Leu196 in Aurora A).

**Table S5:** Comparison of conformational labels with Möbitz’s classification (number of chains)

| Spatial group | Clusters | Möbitz DFGin |  |  |  |  |  |  | Möbitz DFGout |  |  |  |  |
| --- | --- | --- | --- | --- | --- | --- | --- | --- | --- | --- | --- | --- | --- |
|  |  | Active | DFGact<br>Chelixout | FG<br>down | FGdown<br>Chelixout | Gdown | Gdown<br>Chelixout | AuP<br>Met | DFG<br>Flipped | DFGout<br>Type2 | AuP<br>BRAF | AuP<br>FMS | AuP<br>IGF1R |
| DFGin | BLAminus | 1805 | 5 | 1 | 1 | 1 | 7 | - | - | - | - | - | - |
|  | BLAplus | 1 | 13 | 38 | 7 | - | - | - | - | - | - | - | - |
|  | ABAminus | 198 | - | - | - | 69 | 1 | - | - | - | - | - | - |
|  | BLBminus | 5 | - | 5 | 6 | 31 | 49 | - | - | - | - | - | - |
|  | BLBplus | - | 1 | 78 | 125 | - | 48 | 34 | - | - | - | - | - |
|  | BLBtrans | - | - | - | 179 | - | - | - | - | - | - | - | - |
|  | Noise | 43 | 2 | 2 | 16 | 2 | 6 | 1 | 3 | - | - | - | - |
| DFGout | BBAminus | - | - | - | - | - | - | - | - | 73 | 31 | 20 | 1 |
|  | Noise | - | - | - | - | - | - | - | 12 | 2 | 7 | 6 | 4 |
| DFGinter | BABtrans | - | - | - | - | - | - | - | 13 | - | - | - | - |
|  | Noise | - | - | - | - | - | - | - | 6 | - | - | - | - |

**Table S6:**
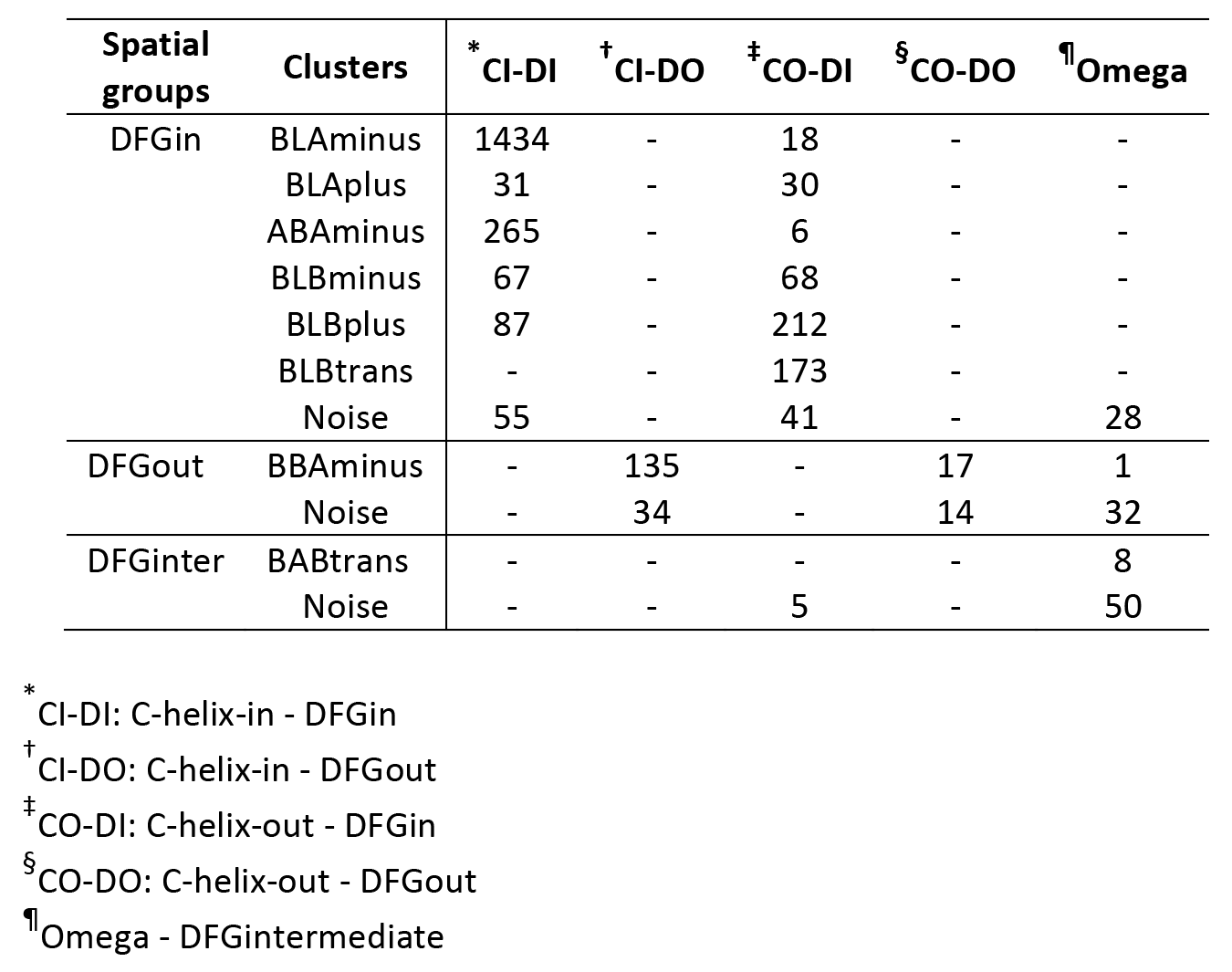
Comparison of conformational labels with Ung et al.’s classification (number of chains)

**Table S7:**
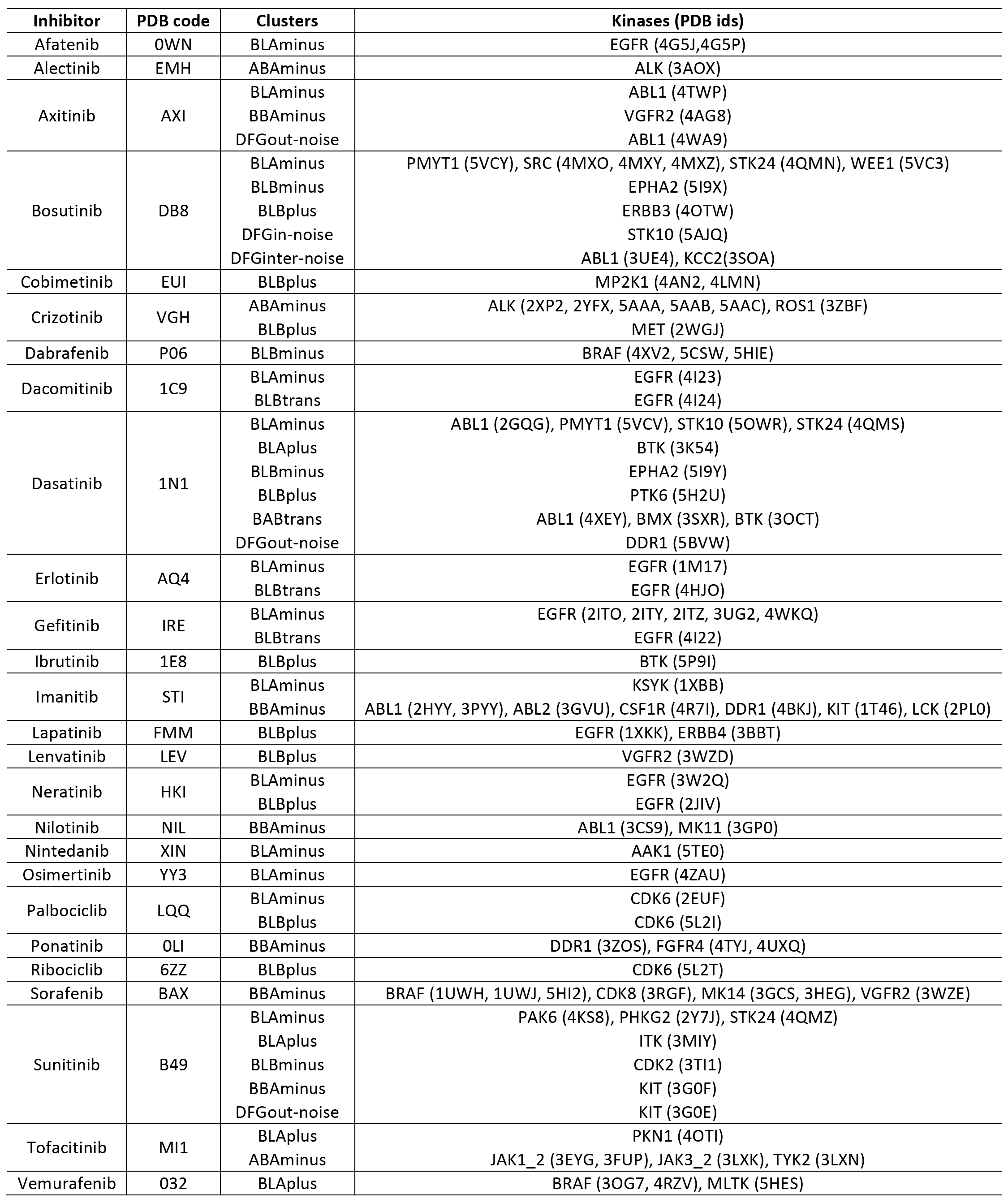
FDA approved inhibitors in complex with structures from different clusters.

**Fig. S1.**
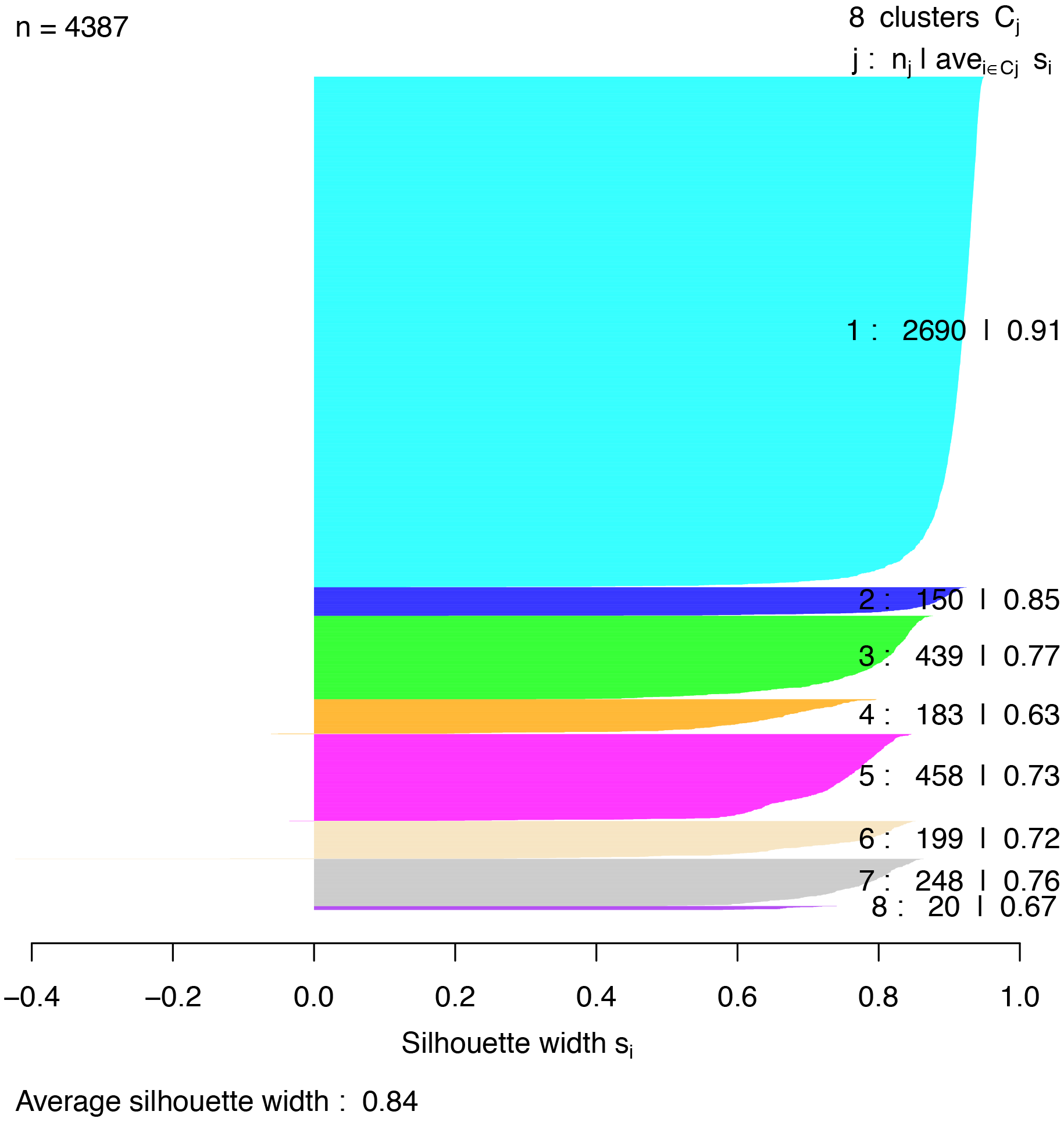
Silhouette scores for all the clusters. Average value over all data points is 0.84.

**Fig. S2.**
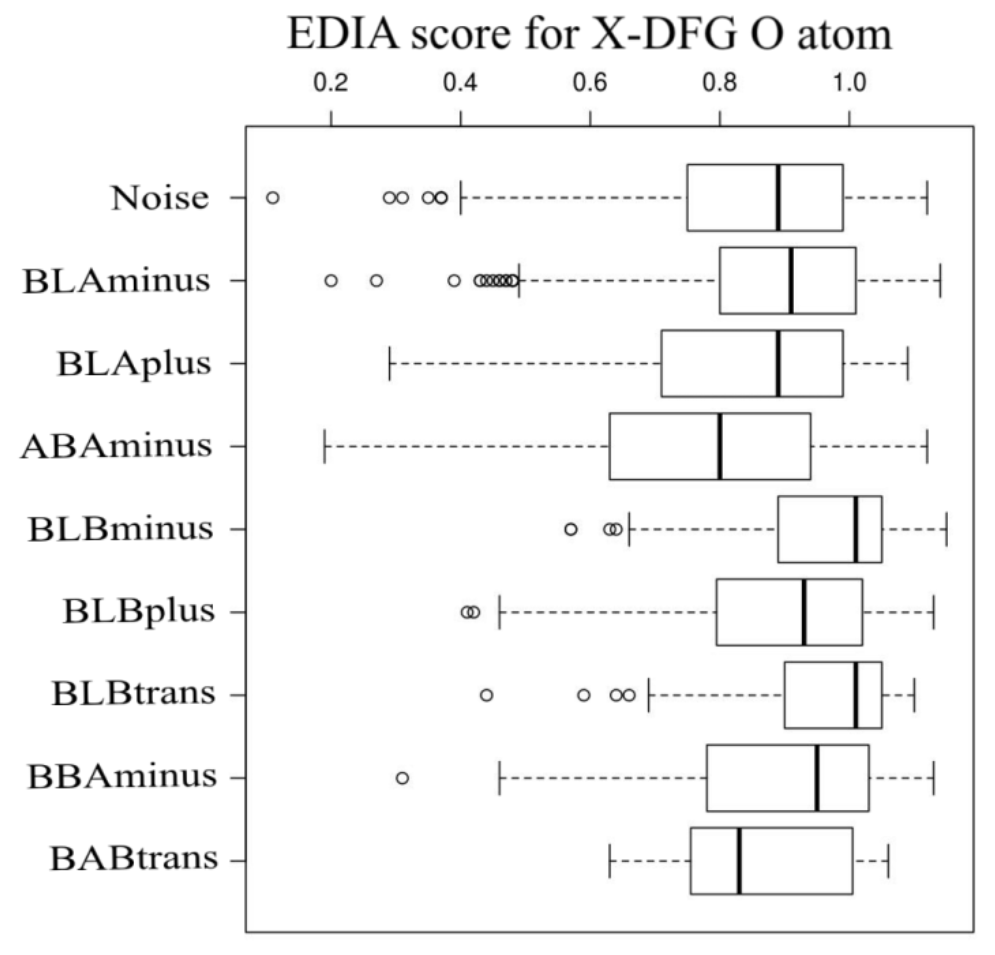
EDIA score of X-DFG O atom across different clusters. An atom is modeled correctly if its score is 0.8 or more.

**Fig. S3.**
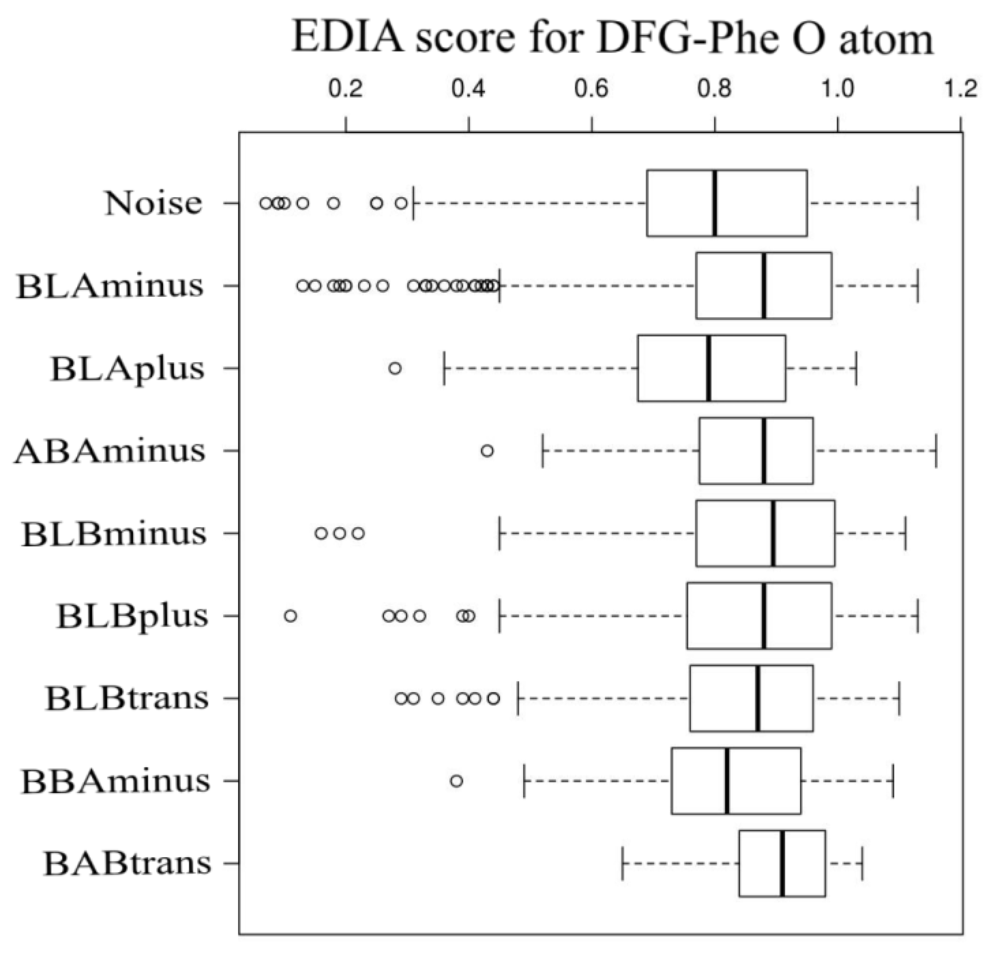
EDIA score for DFG-Phe O atom.

**Fig. S4.**
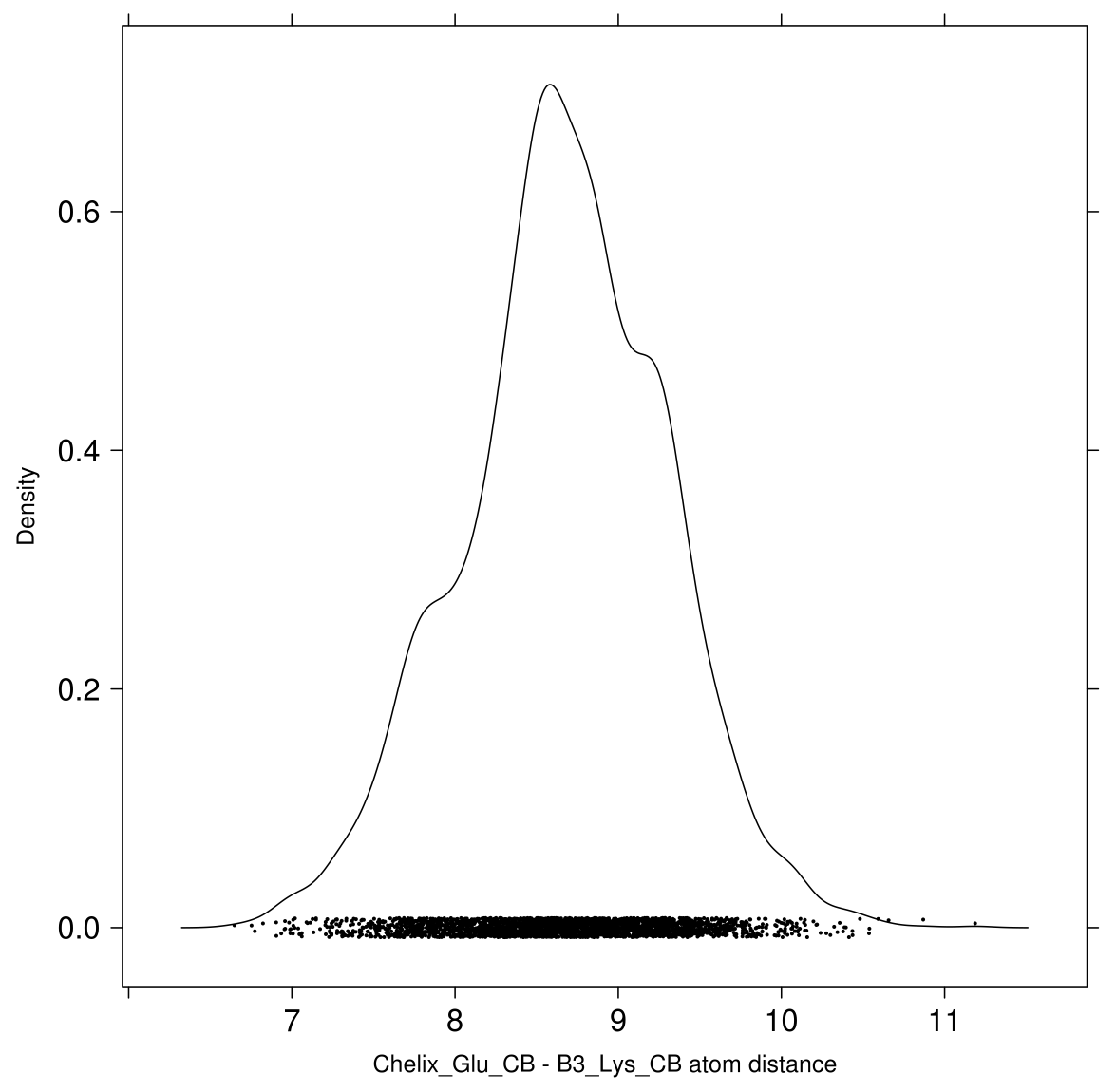
Distribution of distances between C-helix-Glu-CB and β3-Lys-CB atoms in the structures which have an intact salt bridge between these residues (3327 chains, 200 kinases).

**Fig. S5.**
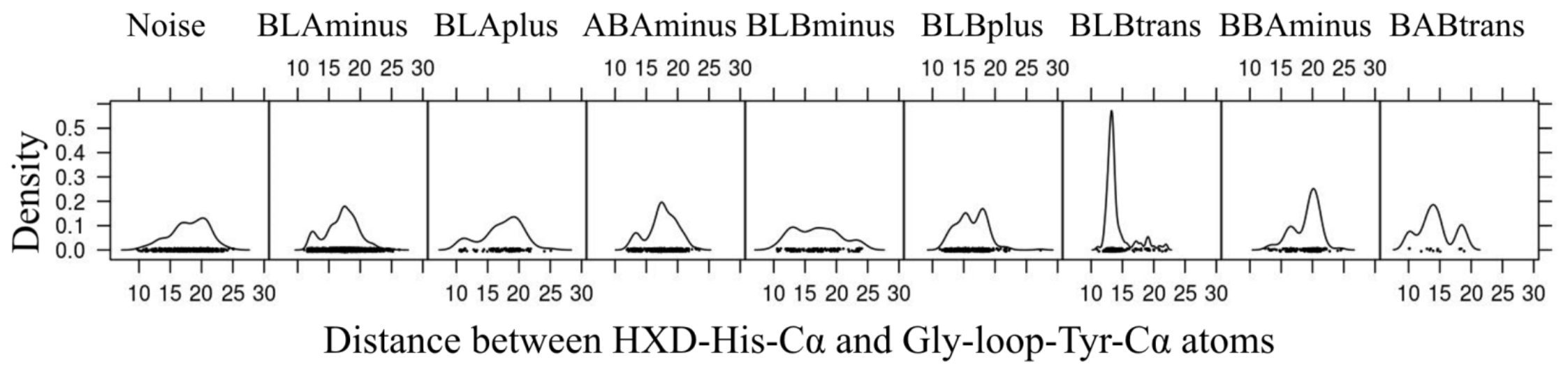
Distribution of Gly-rich loop conformations - distance between HRD-His-Cα and Gly-rich loop-Tyr-Cα atoms (Phe144 in Aurora A kinase).

